# A revised mechanism for how Plasmodium falciparum recruits and exports proteins into its erythrocytic host cell

**DOI:** 10.1101/2021.09.29.462329

**Authors:** Mikha Gabriela, Kathryn M. Matthews, Cas Boshoven, Betty Kouskousis, David L. Steer, Thorey K. Jonsdottir, Hayley E. Bullen, Brad E. Sleebs, Brendan S. Crabb, Tania F. de Koning-Ward, Paul R. Gilson

## Abstract

*Plasmodium falciparum* exports ~10% of its proteome into its host erythrocyte to modify the host cell’s physiology. The *Plasmodium* export element (PEXEL) motif contained within the N-terminus of most exported proteins directs the trafficking of those proteins into the erythrocyte. To reach the host cell, the PEXEL motif of exported proteins are processed by the endoplasmic reticulum (ER) resident aspartyl protease plasmepsin V. Then, following secretion into the parasite-encasing parasitophorous vacuole, the mature exported protein must be unfolded and translocated across the parasitophorous vacuole membrane by the *Plasmodium* translocon of exported proteins (PTEX). PTEX is a protein-conducting channel consisting of the pore-forming protein EXP2, the protein unfoldase HSP101, and structural component PTEX150. The mechanism of how exported proteins are specifically trafficked from the parasite’s ER following PEXEL cleavage to PTEX complexes on the parasitophorous vacuole membrane is currently not understood. Here, we present evidence that EXP2 and PTEX150 form a stable subcomplex that facilitates HSP101 docking. We also demonstrate that HSP101 localises both within the parasitophorous vacuole and within the parasite’s ER throughout the ring and trophozoite stage of the parasite, coinciding with the timeframe of protein export. Interestingly, we found that HSP101 can form specific interactions with model PEXEL proteins in the parasite ER, irrespective of their PEXEL processing status. Collectively, our data suggest that HSP101 recognises and chaperones PEXEL proteins from the ER to the parasitophorous vacuole and given HSP101’s specificity for the EXP2-PTEX150 subcomplex, this provides a mechanism for how exported proteins are specifically targeted to PTEX for translocation into the erythrocyte.

**Author Summary:** *Plasmodium falciparum*, the most lethal species of human malaria parasite, infects erythrocytes and develops within a parasitophorous vacuole. To support rapid parasite growth and immune evasion, the parasite remodels its erythrocyte by exporting a myriad of proteins into the erythrocyte compartment. Parasite proteins destined for export are first imported into the endoplasmic reticulum (ER) and then secreted into the parasitophorous vacuole, where they are translocated across the parasitophorous vacuole membrane into the erythrocyte via a protein-conducting channel called PTEX. A missing link in the story has been how proteins destined for export are specifically guided from the ER to PTEX at the parasitophorous vacuole membrane. In this study, we found that one of the core PTEX components, HSP101, resides within the parasite’s ER, in addition to its PTEX-related location at the parasitophorous vacuole. We also found that ER-located HSP101 can interact transiently with cargo proteins *en route* to the parasitophorous vacuole membrane. Our findings support a model in which HSP101 forms an initial interaction with exported proteins in the ER and then chaperones them to the rest of PTEX at the parasitophorous vacuole membrane for export into the erythrocyte.

## Introduction

Malaria is a parasitic disease responsible for 409,000 deaths in 2019, of which 67% were children under the age of five [1]. The severity and clinical symptoms of malaria are greatly influenced by the ability of the *Plasmodium* parasite to replicate asexually inside its host erythrocyte and extensively renovate it [2]. During each cycle of asexual replication, *Plasmodium falciparum*, the most lethal species of human malaria parasite, invades an erythrocyte and encases itself in an invagination of the host-cell membrane that becomes the parasitophorous vacuole (PV) [3]. The parasite then exports hundreds of effector proteins into its host erythrocyte to radically transform its physicochemical properties. Some notable modifications include the formation of new permeability pathways (NPPs) for plasma nutrient uptake [4–8] and Maurer’s cleft organelles for the transit of exported proteins to the erythrocyte surface [9]. Exported proteins also form knob structures at the erythrocyte surface that display the major virulence protein PfEMP1 on the surface to facilitate the cytoadherence of infected erythrocytes to the microvascular endothelium [10, 11]. All these modifications greatly contribute to chronic parasite infection and the pathological symptoms of malaria [2].

All exported proteins contain one or more transmembrane domains that direct their proteins to the parasite’s endoplasmic reticulum (ER). Most exported proteins also contain a unique (RxLxE/Q/D) motif termed *Plasmodium* export element (PEXEL) or vacuolar transport signal [12, 13] that help feed their proteins into a dedicated trafficking pathway that eventually targets the proteins to specific locations within the host erythrocyte [14]. The remaining subset of exported proteins which lack a PEXEL motif are called PEXEL-negative exported proteins, or PNEPs, and the absence of definitive determinants for export has been a major roadblock for identifying and estimating the exact number of PNEPs [15]. Regardless, it has been estimated that PEXEL proteins and PNEPs collectively enumerate about 10% of the *P. falciparum* proteome [12, 13, 15–19].

Soluble PEXEL proteins are proposed to be post-translationally translocated into the ER lumen via the Sec61-SPC25-plasmepsin V translocon complex, after which the PEXEL motif is cleaved in between the 3^rd^ and 4^th^ residue (RxL^v^ xE/Q/D) by plasmepsin V [18, 20–23]. N-terminal acetylation of the remaining PEXEL motif then occurs, leading to the formation of a mature Ac-xE/Q/D N-terminus on the protein [20]. Post-maturation, the PEXEL protein is secreted into the PV where it is unfolded and translocated across the PVM in an ATP-dependent manner to the host cell [24, 25] by a novel and *Plasmodium*-specific multiprotein complex named *Plasmodium* translocon of exported proteins (PTEX) [26].

PTEX comprises three core components: the pore-forming membrane protein EXP2, the structural component PTEX150, and the protein unfoldase HSP101, which are arranged into a heptamer-heptamer-hexamer conformation, respectively [27–31]. Two distinct conformations of PTEX captured by cryo-EM gave rise to a proposed mechanism of cargo translocation whereupon HSP101, via a ratchet-like movement, threads a cargo polypeptide through the funnel-like membrane pore formed by the N-terminal part of EXP2 [29]. Knocking down or disabling the core components of PTEX leads to loss of protein export and parasite death, highlighting the critical requirement of PTEX and protein export for parasite survival [32–34]. One of the most long-standing questions in this field is how does plasmepsin V cleavage of PEXEL proteins license them for export? Specifically, what factors recognise cleaved PEXEL proteins and escort them to PTEX to the exclusion of non-exported PV resident secreted proteins? It has been previously proposed that ER-resident HSP70 (BIP) and HSP101 may help chaperone PEXEL proteins from the ER to the PV for PTEX mediated export [35, 36]. In support of this role for HSP101, fluorescence microscopy of HSP101 indicates some of the protein is located within the parasite as well as in the PV where EXP2 and PTEX150 almost exclusively reside [37–39].

Here, using conditional knockdown system to deplete HSP101 expression, we found that EXP2 first interacts with PTEX150 to form a stable platform for HSP101 docking. We present biochemical evidence to indicate that a substantial amount of HSP101 resides in the ER and is secreted into the PV during the active protein export period. Lastly, we show that ER-located HSP101 can interact with PEXEL reporter proteins, irrespective of their PEXEL cleavage status. Overall, our data indicates that HSP101 chaperones proteins destined for export from the ER to the PV, whereupon the HSP101/cargo complex docks with the rest of PTEX to facilitate protein translocation into the host erythrocyte compartment.

## Results

### Knockdown of HSP101 blocks protein export and arrests parasite proliferation

To enable functional studies and accurate analysis of HSP101’s subcellular localisation, we generated a transgenic parasite line expressing a triple hemagglutinin (HA) tagged-HSP101 for antibody detection and appended the gene with a *glmS* riboswitch so its expression could be inducibly knocked down [40]. To create the *Pf*HSP101-HA*glmS* line, *P. falciparum* 3D7 parasites were transfected with the pHSP101-HA*glmS* plasmid and were cycled on/off WR99210 until the plasmid had integrated into the HSP101 locus (Fig 1A). PCR analysis of two clonally derived lines using oligonucleotides that distinguish between wildtype and integrant genotypes revealed that integration of the HA*glmS* sequence into the *hsp101* locus had occurred in both clones (Fig 1B). Western blot analysis using anti-HA IgG, further confirmed the presence of HA-tagged HSP101 protein (Fig 1C). Addition of 2.5 mM glucosamine (GlcN) at ring-stage resulted in ~75 % (SD ± 22.3%, n=4) knockdown of HSP101 (Fig 1D), leading to parasites stalling at ring-trophozoite transition the following cycle and parasite death (Fig 1E) as well as a failure to export proteins (Fig 1F). The effects of *glmS-*induced knockdown of PfHSP101 that we observed were consistent with the previously reported phenotypes of the knockdown of *P. berghei* HSP101 [33] and conditional inactivation of HSP101 in *P. falciparum* [32].

**Fig 1.**
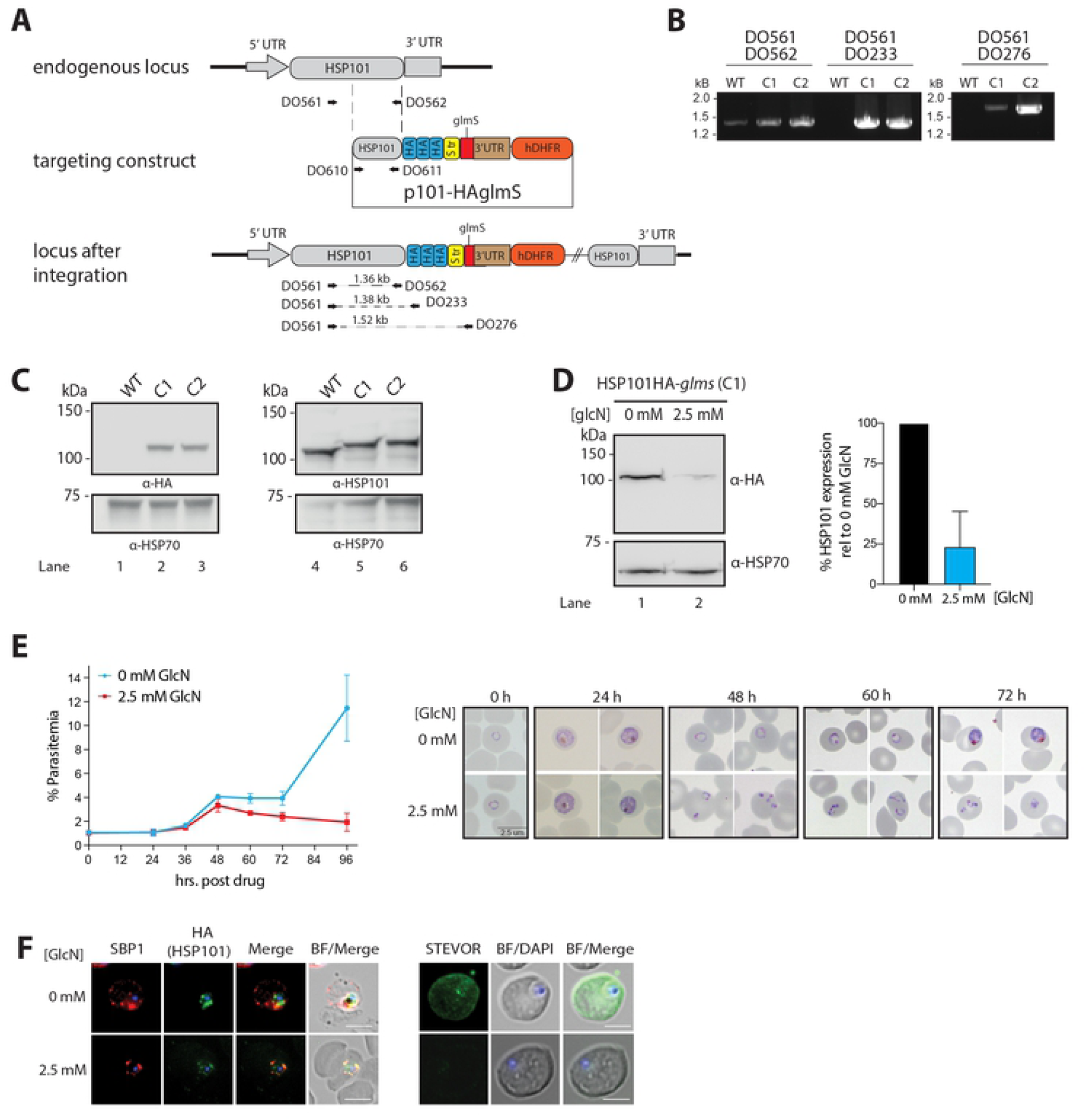
Generation and validation of HSP101-HA*glmS* parasite line and knockdown analysis. (A) The *P. falciparum* HSP101 targeting construct integrated into the endogenous locus by a single crossover recombination event as indicated. Haemagglutinin (HA) and strep II (Str) epitope tags, human DHFR selectable marker (hDHFR), *glmS* ribozyme and untranslated regions (UTR) are shown. Arrows indicate oligonucleotides used in diagnostic PCR analysis and their product sizes. (B) Diagnostic PCR showing the *hsp101* gene contains the integrated sequence. Oligonucleotide pairs shown in (A) were used on genomic DNA prepared from 3D7 wildtype parental parasites (WT) or from two independent clones of drug-resistant parasites obtained after transfection with the targeting construct (C1, C2). DO561 and DO562 recognize the *hsp101* locus and serve as a positive control for the PCR. (C) Western blot analysis showing that the HSP101-HA*glmS* C1 and C2 express the HA epitope tags. Rabbit HSP101 and HSP70-1 antibodies serves as a loading controls. (D) Representative western blot demonstrating HSP101 expression can be knocked down using glucosamine (GlcN). Left panel: Synchronised cultures of ring-stage PfHSP101-HA*glmS* parasites grown in the presence or absence of GlcN for 48 hours were harvested and parasite lysates were probed with anti-HA antibodies to detect HSP101-HA expression. HSP70-1 served as a loading control. Right panel: Densitometry performed on bands observed in western blot using ImageJ to quantitate the level of HSP101 expression levels relative to HSP70-1 in parasite lines grown in the presence or absence of GlcN. Shown is the mean ± SEM (n=3 biological repeats). (E) Parasitemia of cultured PfHSP101-HA*glmS* parasites grown in 0 mM or 2.5 mM GlcN (left panel) and representative Giemsa-stained smears (right panel) shows knockdown of HSP101 leads to the arrest of parasite growth in ring stages, with parasites unable to transition into trophozoite stage. Shown is the mean ± SEM (n=3 biological repeats). (F) Immunofluorescence analysis of ring-stage parasites treated with 2.5 mM GlcN for 48 h and fixed with acetone/methanol and labelled with anti-HA antibodies to detect HSP101-HA, or antibodies against exported proteins SBP1 and STEVOR. This demonstrates that knocking down expression of HSP101-HA blocks the export of SBP1 and STEVOR into the host erythrocyte. Parasite nuclei were stained with DAPI (4’,6-diamidino-2-phenylindole). Scale bar, 5 µm.

### EXP2 and PTEX150 form a stable subcomplex in the absence of HSP101

Upon establishing the inducible HSP101 knockdown line, we first investigated the assembly of the PTEX complex. For this, we employed our HSP101-HA*glmS* line (Fig 2A) along with the previously published PTEX150-HA*glmS* (Fig 2B) and EXP2-HA*glmS* (Fig 2C) parasite lines [33, 34], to enable knockdown of all core PTEX components via GlcN-induced mRNA degradation. Native protein complexes from erythrocytes infected with late ring stage parasites that had been treated +/− 2.5 mM GlcN for one cell cycle were fractionated by blue native PAGE (BN-PAGE) and analysed via western blotting. In the absence of GlcN treatment, a >1236 kDa band, consisting of all three core components of PTEX [27, 28], was common in all parasite lines (Fig 2A, 2B, and 2C, odd numbered lanes). In the same lanes, PTEX150 and EXP2 formed smaller oligomers of ~500 kDa and ~700 kDa, respectively, whilst HSP101 produced oligomers of >720 kDa, consistent with what has been previously described (Fig 2A) [27, 28, 41]. Strikingly, we observed that following the knockdown of the HSP101 component, PTEX150 and EXP2 were still present in the >1236 kDa, although this complex was now devoid of most HSP101 (Fig 2A, lanes 4 and 6). Knockdown of HSP101 also revealed that anti-HSP101 antibody cross reacted with ~1000 and ~200 kDa bands that were not recognised by the anti-HA antibody (Fig 2A, lane 2 asterisks and S1A Fig). In contrast, individual knockdown of both PTEX150 and EXP2 resulted in the loss of the >1236 kDa complex (Fig 2B, lanes 8 and 12; Fig 2C, lanes 14 and 16). This data suggests that PTEX150 and EXP2 are still able to associate and form a large subcomplex that is not dependent on the HSP101 association (Fig 2D).

**Fig 2.**
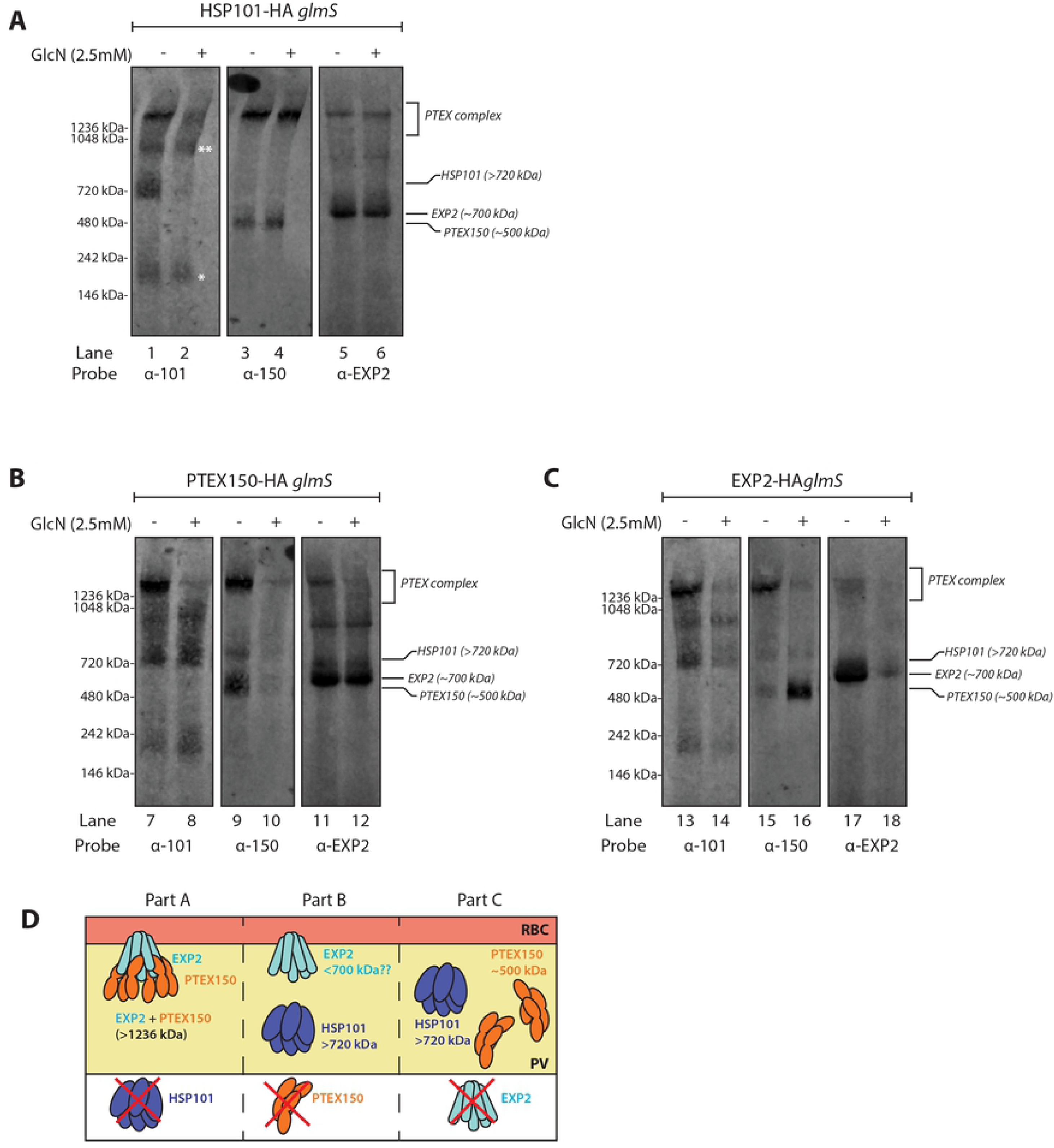
Knockdown of HSP101 does not disrupt the stability of the EXP2/PTEX150 subcomplex. (A), (B), and (C) BN-PAGE analysis of PTEX-HA*glmS* parasite lines following protein knockdown enables the effects on PTEX complex formation to be examined. Saponin-lysed pellets were lysed with 1% digitonin, separated on 4-12% NativePAGE gel, and analysed via western blotting using PTEX-specific antibodies. Knockdown of EXP2 and PTEX150 prevented the formation of a >1236 kDa PTEX subcomplex. Knockdown of HSP101 in contrast, did not disrupt the ability of PTEX150 and EXP2 to form a >1236kDa complex free of HSP101. * and ** are two non-specific bands observed with rabbit anti-HSP101 antibody (D) Schematic summarising the impact of knocking down individual PTEX components with regards to complex formation.

One could anticipate that the PTEX150-EXP2 subcomplex would have a lower molecular weight relative to the full translocon due to the loss of HSP101. Interestingly, this does not seem to be the case as we did not observe a reduction in the size of the >1236kDa EXP2 and PTEX150 subcomplex after HSP101 knockdown (Fig 2A, lanes 3-6). However, assuming that PTEX150 and EXP2 retained the heptameric-heptameric arrangement of the full PTEX complex in the absence of HSP101 [29], the resulting tetradecameric subcomplex of PTEX150-EXP2 would have a theoretical size of approximately ~1300 kDa, which is still too large for the BN-PAGE system to resolve [29].

Notably, the loss of the >1236 kDa complex upon EXP2 knockdown caused a remarkable accumulation of PTEX150 ~500 kDa oligomer (Fig 2C, lane 16) that did not label with anti-HSP101 antibody (Fig 2C, lane 14). This observation suggests that in the absence of EXP2, PTEX150 alone cannot associate with HSP101, and forms a different oligomeric state. There was no equivalent increase of the EXP2 ~700 kDa oligomer upon destabilisation of the >1236 kDa complex by knockdown of PTEX150 (Fig 2B, lane 12) or HSP101 (Fig 2A, lane 6). The EXP2 knockdown, however, clearly reduced the level of the ~700 kDa oligomer suggesting that perhaps the EXP2 oligomer is not a part of PTEX complex (Fig 2C, lane 18). The ~700kDa EXP2 species may therefore represent a dedicated non-translocon EXP2 pool which serves other functions, such as forming the PVM nutrient channel [42]. As observed with the EXP2 knockdown, we did not see the appearance of subcomplexes consisting of EXP2 and HSP101 upon PTEX150 knockdown, suggesting that EXP2 similarly is unable to interact with HSP101 in the absence of PTEX150.

To validate this finding, we performed co-immunoprecipitations to examine direct interactions between each of the core PTEX components following the knockdown of each subunit (S1B and S1C Fig). As expected, knockdown of HSP101-HA*glmS* with 2.5 mM GlcN followed by immunoprecipitation of PTEX150 from parasite lysates revealed a non-significant change (p=0.9319, n=3) in PTEX150 interaction with EXP2, despite little HSP101 being present (S1B Fig, lanes 3 and 4, S1C Fig). In contrast, knockdown of EXP2 resulted in less interaction between PTEX150 and HSP101 (p=0.0036, n=3, S1B Fig, lanes 7 and 8, S1C Fig). Similarly, there was a trend for reduction in EXP2 and HSP101 interaction upon knockdown of PTEX150 (S1B Fig, lanes 11 and 12, S1C Fig), although notably, this was not statistically significant (p=0.3239, n=3), S1C Fig), presumably due to inefficient PTEX150 knockdown (~20% reduction in protein expression). Nevertheless, the co-immunoprecipitations support the BN-PAGE data and suggest that following redcution of HSP101, PTEX150 can still engage with EXP2 to form the large >1236 kDa subcomplex. Collectively, our data indicate that PTEX assembles in two broad steps where EXP2 and PTEX150 first assemble to form a stable subcomplex, which then enables HSP101 docking to form the fully functional translocon.

### HSP101 localises to the parasitophorous vacuole and within the parasite

We have previously reported that HA-tagged HSP101, displays dual localisation in the PV and inside the parasite [38]. Interestingly, an independent study also showed that the 3xFLAG-tagged HSP101 displayed a similar localisation pattern [39]. We therefore decided to investigate this phenomena in more detail. We first confirmed the dual localisation of HSP101 in our HSP101-HA*glmS* line by performing immunofluorescence assays (IFAs) on paraformaldehyde/glutaraldehyde fixed cells. Anti-HA staining indeed showed labelling at two main sites; the parasite’s periphery, and the perinuclear area inside the parasite (Fig 3A, panels 1 and 2). This staining pattern can be observed throughout ring to early-mid trophozoite stage parasites, coinciding with the duration of active protein export [28, 32]. To confirm that this staining is an accurate representation of HSP101 localisation rather than non-specific staining of our anti-HA antibody, PTEX150-HA*glmS* and EXP2-HA*glmS* parasites were also labelled using the same anti-HA antibody and imaged under the same conditions. In contrast to the anti-HA labelling that localised HSP101-HA at the parasite’s periphery and internally (Fig 3A, panels 1 and 2), both PTEX150-HA and EXP2-HA localised solely at the parasite’s periphery, consistent with previous observations (Fig 3A, panels 3-6) [26], confirming that the internal anti-HA labelling observed in the HSP101-HA*glmS* parasite line was specific.

**Fig 3.**
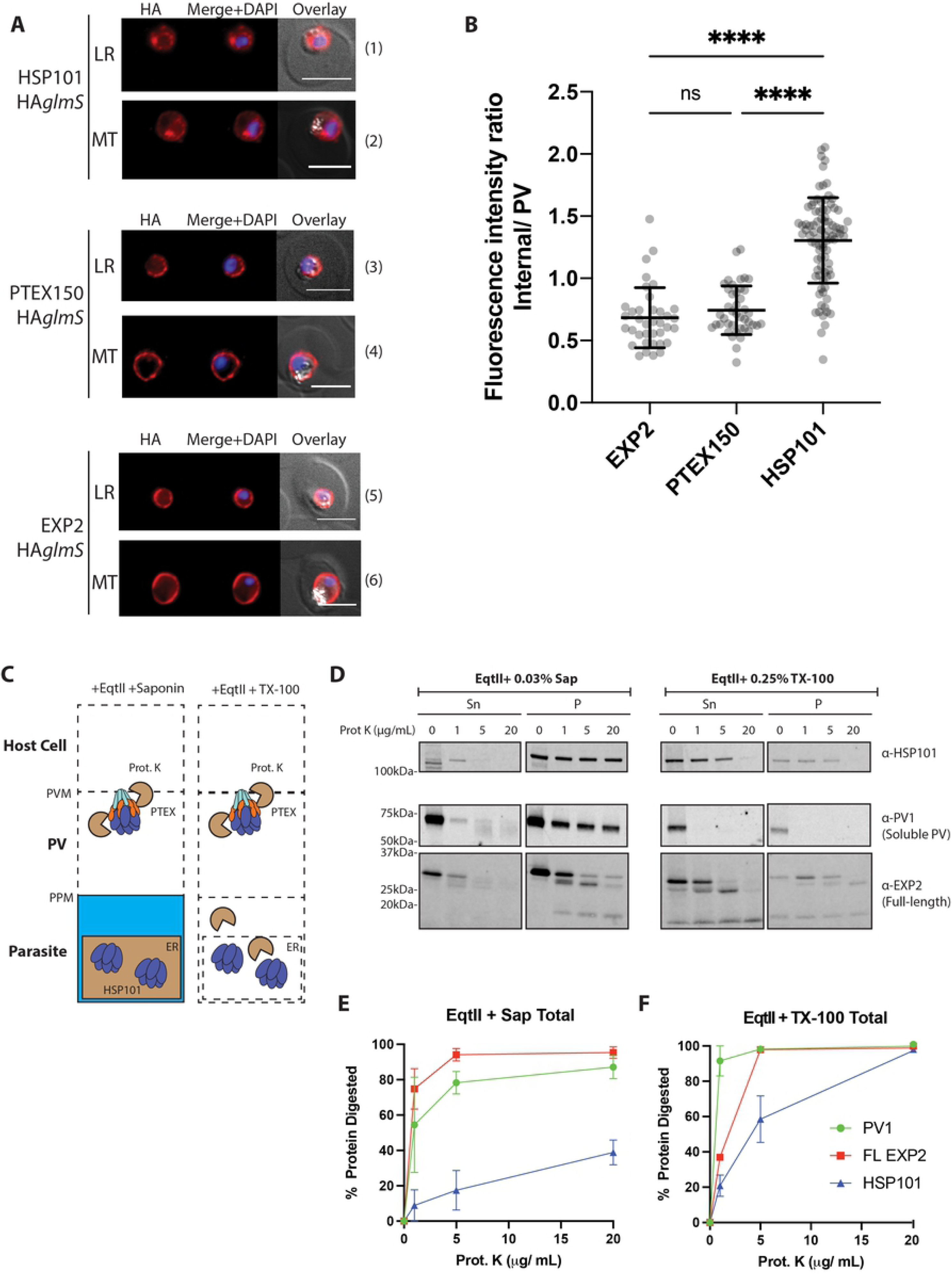
HSP101 exhibits distinct dual localisation inside the parasite and in the parasitophorous vacuole. (**A)** Representative IFA images of parasites expressing HSP101-HA, EXP2-HA, or PTEX150-HA. Parasites were labelled with anti-HA IgG and nuclei were stained with DAPI. LR, Late ring stage. MT, Mid trophozoite stage. Scale bar, 5 µm. (B) Graph of the ratios of internal parasite signal over PV signal for each cell line indicates HSP101-HA exhibits a comparatively greater internal signal than the other PTEX proteins. Data were pooled from at least two biological replicates, with >20 cells measured for each line. Statistical significances were determined using an ordinary one-way ANOVA. ****, p-value<0.0001). (C) Schematic of the proteinase K (Prot. K) protection assay to validate cellular localisation of HSP101. EqtII, equinatoxin II; TX-100, Triton X-100 (D) Western blots of permeabilised mid-trophozoite stage parasites digested with increasing concentrations (0-20 µg/mL) of proteinase K. Sn, Supernatant. P, Pellet. (E) Densitometry analysis of the western blot in (D) showing the total protein digested with Prot. K in saponin (Sap) or TX-100-treated parasites from the combined signals of the Sn and P fractions. Percentage normalised to the total band intensity (Sn and P fractions) of samples treated with 0 µg/mL Prot. K. The data indicate that compared to known PV markers EXP2 and PV1, more HSP101 is resistant to proteolytic degradation in saponin suggesting a large pool of HSP101 resides inside the parasite. Data were extracted from two independent biological replicates. Plotted data represents the mean ±SEM.

To quantify the relative proportion of cells exhibiting dual localisation of HSP101-HA in our parasite population, the intensity of internal and peripheral fluorescence signals was measured separately for more than 20 cells per line and expressed as an internal/PV ratio. A higher internal/PV ratio would therefore indicate that the measured cell has more internal signal. Indeed, HSP101-HA staining displayed a higher overall internal/PV ratio relative to EXP2-HA and PTEX150-HA, suggesting that HSP101-HA localises within the parasite in a significant number of cells in the population (Fig 3B).

We also sought to validate this finding using a biochemical approach. For this, we utilised a proteinase K protection assay to assess the susceptibility of HSP101 and EXP2 to proteolytic degradation upon erythrocyte and PVM permeabilisation by equinatoxin II and saponin (Fig 3C) [43]. We have previously used this assay to show that the C-terminal region of EXP2 resides in the PV lumen and can be readily degraded by proteinase K upon PVM permeabilization to leave only the N-terminal transmembrane domain intact [30]. We assumed that if an internal parasite pool of HSP101 exists, it would be protected against protease degradation at a concentration sufficient to fully degrade the C-terminal part of EXP2. Quantitative proteolysis of full-length EXP2 after equinatoxin II and saponin treatment indicated that in the supernatant and pellet fractions, EXP2 was degraded into smaller fragments that when combined, resulted in >90% degradation in the presence of 20 µg/mL proteinase K (Fig 3D and 3E). In comparison, treatment of HSP101 with 20 µg/mL proteinase K only degraded 40% of the total full-length protein for combined soluble and pellet fractions, with the degraded and resistant proportions likely representing vacuolar and internal parasite HSP101, respectively [26, 27].

To control for PVM permeabilization, we used a parasite line expressing a HA epitope-tagged version of the PV-resident protein PV1 [44]. About half of PV1 was soluble suggestive of vacuolar localisation, and half was in the pellet fraction indicative of an internal parasite pool as the protein is being actively made during this period. As expected, the soluble vacuolar fraction of PV1 was highly susceptible to proteinase K treatment, indicating that the PVM had been permeabilised (Fig 3D and 3E). The remaining pellet fraction of PV1 was partially degraded with 20 µg/mL proteinase K consistent with this fraction presumably containing both newly synthesised internal PV1 and PVM-associated protein since some PV1 is known to bind to PVM-bound PTEX and EPIC [44]. Overall, 80% of PV1 was degraded at the highest PK concentration indicating the PVM was permeabilised (Fig 3D and 3E).

When parasites were treated with equinatoxin II and 0.25% Triton X-100 to disrupt all parasite membranes, EXP2, HSP101, and PV1 were almost completely degraded by treatment of 20 µg/mL proteinase K, indicating the three proteins were susceptible to proteinase K treatment when completely solubilised (Fig 3E). Thus, both microscopy and biochemical data support the presence of an internal pool of HSP101.

### The internal pool of HSP101 resides within the endoplasmic reticulum of the parasite and is secreted into the parasitophorous vacuole

Having confirmed the presence of an internal pool of HSP101 in our parasites, we next sought to determine HSP101’s subcellular localisation. By IFA, the perinuclear HSP101 signal was reminiscent of ER localisation. Indeed, *P. berghei* HSP101 has also been reported to localise to the ER in ring and trophozoite stages [37]. To test whether the internal HSP101 we observed in *P. falciparum* also localised to the ER, IFAs were first performed on tightly synchronised HSP101-HA*glmS* parasites at the late ring stage, where the parasite cell size is sufficiently large to clearly visualise dual localisation of HSP101 (Fig 3A, panels 1 and 2). As expected, immunolabelling with anti-HA and ER-resident calcium-binding protein PfERC showed co-localisation of the internal pool of HSP101-HA with PfERC, indicative of ER-localisation of PfHSP101 (Fig 4A).

**Fig 4.**
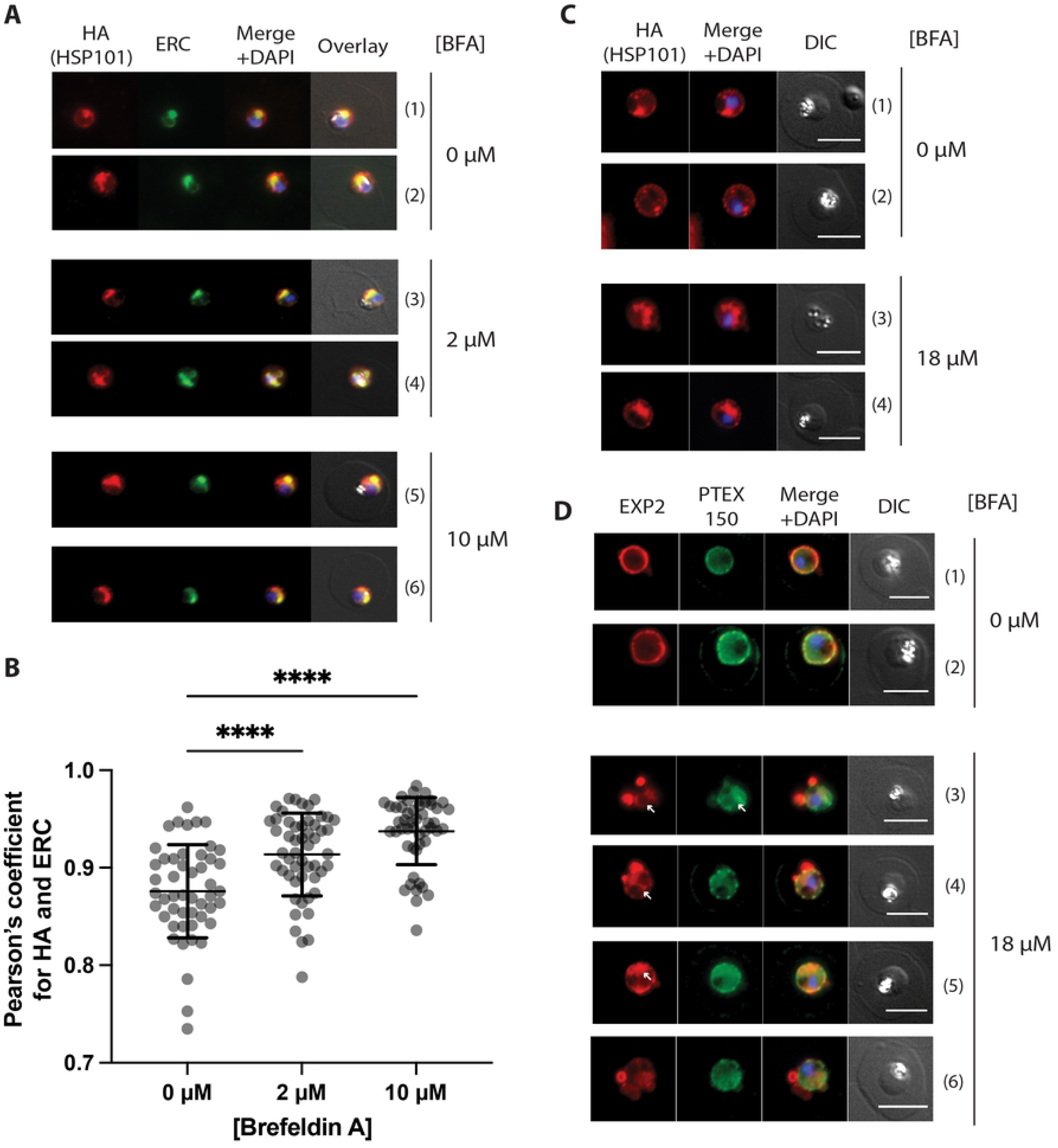
Brefeldin A treatment induces internalisation of HSP101. (A) Representative IFA images of HSP101-HA*glmS* parasites treated with 0, 2, and 10 µM Brefeldin A (BFA) for 24 hours starting from the ring stage. Cells were fixed and stained with anti-HA (HSP101) and anti-ERC antibodies (ER marker). (B) Quantification of the degree of co-localisation (Pearson’s coefficient) between HA and ERC fluorescence signal indicates that HSP101-HA becomes progressively more trapped within the ER at higher BFA concentrations. The Z-stack images of at least 50 cells were analysed for each treatment. Statistical significances were determined using ordinary one-way ANOVA. ****, p-value<0.0001. (C) and (D) Older parasites (16-20 hours post invasion) were treated with 0 and 18 µM BFA for 5 hours and stained with (C), anti-HA (HSP101) or (D), anti-EXP2 and anti-PTEX150. The images indicate that HSP101 becomes more noticeably trapped within the parasite and depleted from the PV than EXP2 and PTEX150 although there is some accumulation of the latter two proteins within parasite after BFA treatment (white arrow, perinuclear staining of EXP2 and PTEX150). Scale bar, 5 µm. DIC, differential interference contrast.

Given the ER-localisation of HSP101, we next asked whether this pool of HSP101 was destined to be trafficked to the PV. To investigate this, we first treated early ring stage HSP101-HA*glmS* parasites (0-4 hours post infection) with an increasing concentration of COPII inhibitor Brefeldin A (BFA) to inhibit ER-dependent protein secretion. Parasites were then fixed and labelled with anti-HA and PfERC antibodies 20 hours post treatment, when they reached late ring-early trophozoite stage. As expected, there was a dose-dependent accumulation of HSP101 inside the parasite’s ER as measured by increased co-localisation with anti-PfERC signal by IFA. At 2 and 10 µM BFA concentration, the parasite’s ER increasingly collapsed into a single HSP101 and ERC staining body (Fig 4A, panels 3-6). This accumulation appeared to also coincide with the disappearance of the anti-HA staining at the parasite’s periphery resulting in greater co-localisation of HSP101-HA and ERC (Fig 4B).

It is possible that the ER-located HSP101 represents newly synthesised HSP101 which was recently shown to occur up to 20-24 hours post infection [42, 45]. BFA treatment commencing at the early ring stage overlaps with the period of new HSP101 transcription and protein synthesis and results in retention of the newly synthesised HSP101 in the ER and subsequent depletion of HSP101 in the PV. However, this could not explain why we still sometimes observed ER-localised HSP101 during the mid to late trophozoite stage (>24 hours post infection, Fig 3A, panel 2).

To minimise the confounding effect of newly synthesised protein, we performed the BFA trapping experiment on mid stage trophozoites (24-28 hours post infection) in which there was little new HSP101 being synthesised [42]. Here, parasites were treated with a higher concentration of BFA (18 µM) but for a shorter period (5 hours) to enable visualisation of parasite cells prior to the parasites reaching schizogony. With anti-HA labelling, increased accumulation of HSP101 was still observed in the ER, as well as the disappearance of HSP101 signal from the PV (Fig 4C, panels 1 and 2 versus 3 and 4). Anti-EXP2 and PTEX150 labelling was also performed to gauge the impact of BFA treatment on the localisation of the remaining PTEX components during mid trophozoite stage. Although the transcription profile of PTEX150 matches that of HSP101, BFA-induced ER retention was much less pronounced for PTEX150 and in most cases, PTEX150 remained localised predominantly at the parasite’s periphery (Fig 4D). On occasion, a faint signal of PTEX150 inside the parasite cell can be observed (Fig 4D, panels 3 and 4) that could represent a small amount of trapping of newly synthesised protein. In contrast, EXP2 labelling displayed perinuclear staining more visibly in BFA-treated parasites (Fig 4D, panels 3-5, white arrows), likely because synthesis of new EXP2 peaks during the trophozoite stage [26, 42]. EXP2 staining could also be observed around the parasite’ periphery, but sometimes in the form of PTEX150-free blebs (Fig 4D), perhaps due to the stress caused by failure to secrete proteins important for maintaining proper PVM organisation, or the trapping of cargo proteins [38, 42]. These EXP2 blebs were often not co-localised with PTEX150 (Fig 4D, panel 6), similar to what was previously observed [38, 39]. Overall, the data collectively suggests that the ER-resident pool of HSP101 cannot be sufficiently explained by secretion of newly synthesised HSP101 protein alone since PTEX150, although transcribed at the same time as HSP101, had mostly been secreted into the PV [27, 42]. As only low levels of newly synthesized HSP101 are expressed at this stage, BFA induced PV depletion and ER-trapping may indicate that there is a pool of HSP101 that cycles between the PV and ER compartments [36].

### HSP101 co-localises with a reporter PEXEL protein in the endoplasmic reticulum

It has been previously proposed that HSP101 might chaperone exported proteins from the ER into the PV [35, 36]. The internal localisation of HSP101 and the assembly model of PTEX we have proposed here are consistent with this model. We therefore investigated the possible involvement of HSP101 in the early protein export pathway in the ER where we first sought to determine whether HSP101 in the ER co-localises with proteins destined for export. For this, we generated a reporter construct called Hyp1-Nluc-mDHFR-3xFLAG. This construct contains the first 113 amino acids of Hyp1 (PF3D7_0113300) including 52 amino acids downstream of the plasmepsin V cleavage site [46]. The reporter used here was modified from an earlier version [46] to contain a murine dihydrofolate reductase (mDHFR) domain to stabilise the Hyp1-Nluc cargo within PTEX and a triple FLAG tag [47] to facilitate downstream purification of the protein. Reporter constructs were transfected either to the HSP101-HA*glmS* or HSP101-HA parasite lines [26] and as anticipated, parasite IFAs demonstrated that Hyp1-Nluc-mDHFR-3xFLAG was efficiently exported into the host-cell when expressed in early to late trophozoite parasites under the *P. berghei* EF1a promoter (Fig 5A, panels 1 and 3). Treatment of parasites with the anti-folate compound WR99210, resulted in a stabilised mDHFR domain and trapping of the reporter protein within the parasite and with EXP2 in the PV (Fig 5A, panel 2).

**Fig 5.**
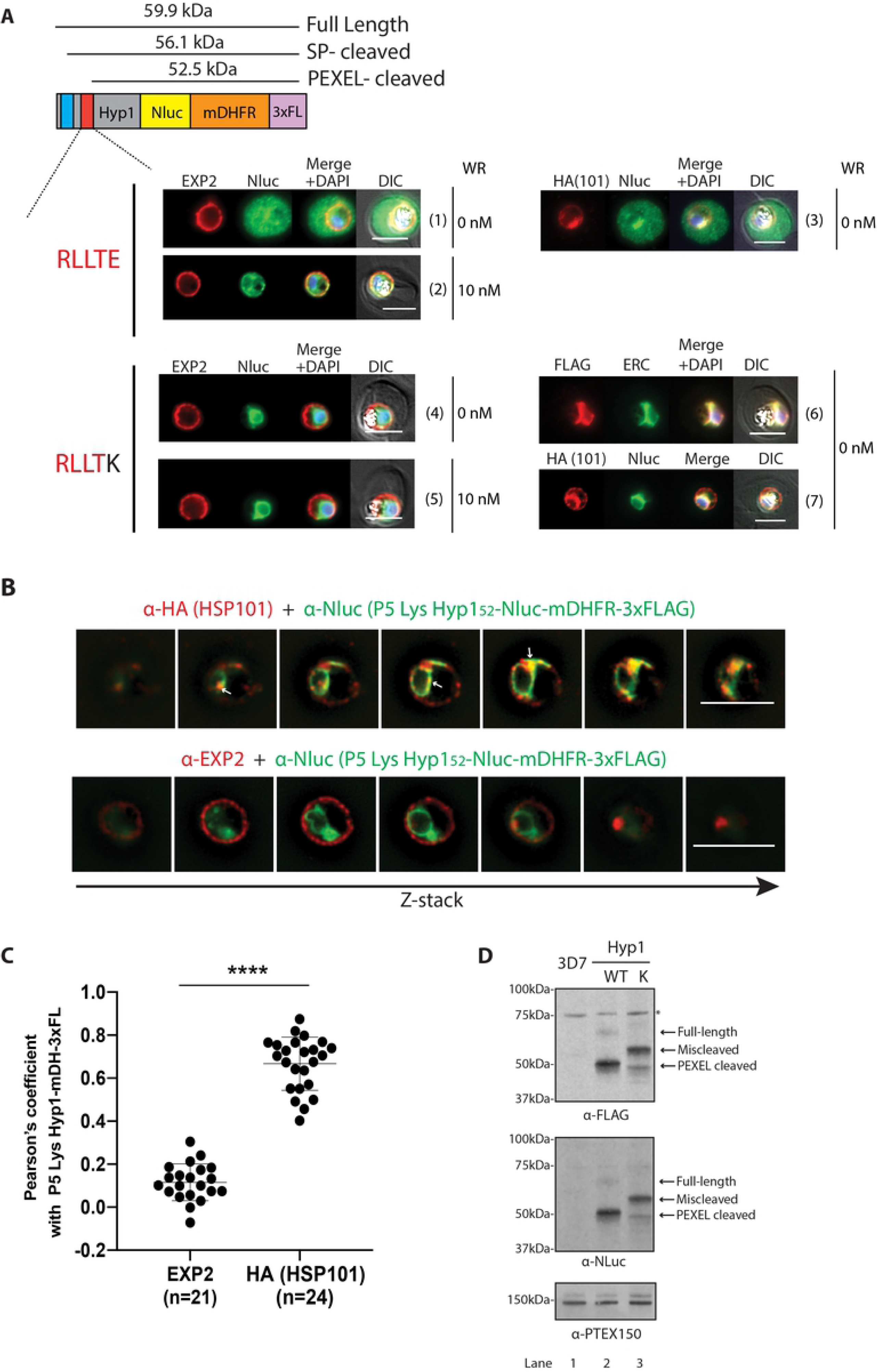
Intra-parasite HSP101 co-localises with P5 Lys Hyp1-Nluc-mDHFR-3xFLAG reporter construct in the endoplasmic reticulum of the parasite. (A) Representative IFA images (n=3 independent replicates) of parasites expressing Hyp1-Nluc-mDHFR-3xFLAG constructs with WT P5 glutamate reporter exported into the erythrocyte and the mutant P5 Lys Hyp1-Nluc-mDHFR-3xFLAG reporter trapped in the parasite ER. To visualise translocon components, cells were probed with anti-EXP2 and anti-HA (to visualise HA-tagged HSP101). Anti-PfERC was used to visualise the parasite’s ER. Hyp1-Nluc-mDHFR-3xFLAG reporter proteins were localised using either anti-Nluc or anti-FLAG antibodies. The red bar in the schematic picture of the construct indicates the PEXEL motif and variations thereof. The blue bar indicates the transmembrane signal peptide. Scale bars, 5 μm. DIC, Differential Interference Contrast. DAPI was used to stain parasite nuclei. (B) Representative Z-stacks of 3D-SIM images of the HA-tagged HSP101 parasite line expressing P5 Lys Hyp1-Nluc-mDHFR-3xFLAG and probed with either anti-HA for HSP101 (top, red) or anti-EXP2 (bottom, red), and anti-Nluc (green) to probe for the cargo. Scale bars represent 5 μm. (C) Degree of co-localisation between individual PTEX components to the P5 Lys Hyp1-Nluc-mDHFR-3xFLAG was calculated by measuring Pearson’s coefficients of the merged Z-stack images of >20 cells. Statistical significances were determined using an unpaired t-test with Welch’s correction. ****, p-value<0.0001). (D) Representative Western blot (n>3) of lysates made from mid-stage trophozoites expressing WT and P5 Lys Hyp1-Nluc-mDHFR-3xFLAG constructs showing full-length and cleaved forms of the WT Hyp1-Nluc-mDHFR-3xFLAG reporter. The P5 Lys Hyp1-Nluc-mDHFR-3xFLAG reporter appears to be miscleaved after the signal peptide (58.1 ± 1.2 kDa, n=10). The blot was probed with anti-PTEX150 antibodies as a loading control.

To test whether HSP101 recognises the Ac-xE/D/Q N-terminus generated after cleavage of the PEXEL motif in the ER, we created a single Glu to Lys charge reversal mutation of the conserved 5^th^ residue of the Hyp1 PEXEL motif as this type of radical mutation has been previously reported to block export [48]. The resulting P5 Lys Hyp1-Nluc-mDHFR-3xFLAG reporter protein failed to be exported as anticipated, but interestingly appeared to co-localise with ERC in the ER regardless of WR99210 treatment (Fig 5A, panels 4-6). We used this ER-trapped construct and assessed its co-localisation with the ER-located HSP101, which revealed a qualitatively high degree of co-localisation around the perinuclear area of the parasite (Fig 5A, panel 7).

To further improve the resolution of subcellular proteins, we imaged HSP101-HA parasites expressing P5 Lys Hyp1-Nluc-mDHFR-3xFLAG using 3D structured illumination microscopy (3D-SIM). Co-labelling of paraformaldehyde/glutaraldehyde-fixed cells with anti-HA and Nluc IgGs revealed partial co-localisation between the P5 Lys Hyp1-Nluc-mDHFR-3xFLAG and the internal pool of HSP101-HA in the perinuclear region and in close apposition to the periphery of the parasite (Fig 5B top panel). But whilst P5 Lys Hyp1-Nluc-mDHFR-3xFLAG signal was distributed evenly (perhaps due to saturation of signal caused by accumulation of the protein in the ER), internal HSP101 sometimes formed small puncta or concentrated to a certain area (Fig 5B, white arrows) that overlapped with P5 Lys Hyp1-Nluc-mDHFR-3xFLAG throughout the z-stack. In contrast, co-labelling of anti-EXP2 with anti-Nluc antibodies revealed almost no co-localisation, consistent with EXP2 residing only at the PVM (Fig 5B, lower panel). Pearson’s coefficients of anti-HA (HSP101) and anti-Nluc co-labelling on maximum projection images yielded an average value of 0.67 (SEM ±0.02, n=24, Fig 5C), which is significantly higher than the co-localisation value for anti-Nluc and anti-EXP2 (0.12 ±0.02, n=21) (Fig 5C). This data suggests that the ER-localised HSP101 and the PEXEL reporter protein localise near one another in the ER.

To understand why the P5 Lys mutation resulted in ER-trapping, protein extract from parasites expressing the WT or P5 Lys Hyp1-Nluc-mDHFR-3xFLAG were separated on SDS-PAGE and analysed by western blot using anti-FLAG and anti-Nluc antibodies. WT Hyp1-Nluc-mDHFR-3xFLAG protein migrated at a predicted size of 50 kDa, consistent with efficient plasmepsin V cleavage within the PEXEL site, although a small amount of larger full-length protein could be observed (Fig 5D). In contrast, P5 Lys Hyp1-Nluc-mDHFR-3xFLAG runs predominantly at a higher molecular weight (~56 kDa), indicative of miscleaved protein, followed by a small proportion of PEXEL cleaved protein (87% and 13% (SD ± 5.3%, n=6), respectively, Fig 5D). To determine if plasmepsin V was able to cleave the mutant reporter, a synthetic Hyp1 PEXEL peptide containing the P5 Lys mutation was treated with recombinant *P. vivax* plasmepsin V since the *P. falciparum* protease is more difficult to express (S2 Fig)[49] Relative to the WT Hyp1 peptide, cleavage of the P5 Lys reporter was reduced >90%, which is consistent with the proportion of miscleaved protein observed in western blots. As the miscleaved 56 kDa P5 Lys Hyp1-Nluc-mDHFR-3xFLAG band is smaller than the full-length band it might be processed upstream of the PEXEL motif, possibly by signal peptidase (Fig 5D). Mass spectrometry of immunoprecipitated forms of both Hyp1 reporter proteins indicated peptide coverage was only present downstream of the PEXEL cleavage site for the WT protein whereas peptides corresponding to the region upstream of the PEXEL were detected with P5 Lys protein (S3 Fig). This data, together with the *in vitro* plasmepsin V cleavage assay, suggests that the PEXEL processing of the P5 Lys Hyp1-Nluc-mDHFR-3xFLAG is significantly reduced, leading to the formation of higher molecular weight protein that has its PEXEL motif intact (S3C Fig). The implication of this observation will be discussed later.

Overall, these results indicate that mutation of the P5 Hyp1 residue to Lys blocks efficient cleavage by plasmepsin V and results in nearly complete trapping of the reporter protein in the ER. Furthermore, a substantial amount of HSP101 co-localises with P5 Lys Hyp1-Nluc-mDHFR-3xFLAG in the parasite and not in the PV.

### ER-resident HSP101 interacts with Hyp1 reporter proteins independent of PEXEL processing

We next performed co-immunoprecipitations to investigate if ER-localised HSP101 forms specific interactions with WT and P5 Lys Hyp1-Nluc-mDHFR-3xFLAG. P5 Lys Hyp1-Nluc-mDHFR’s exclusive ER-localisation would allow us to assess its interaction with internal HSP101 only. In addition, the fact that the PEXEL motif of P5 Lys Hyp1-Nluc-mDHFR-3xFLAG is intact (Fig 5D and S3 Fig), enabled us to determine if PEXEL processing is crucial for the interaction with ER-localised HSP101. For this experiment, HSP101-HA*glmS* parasites expressing the Hyp1 reporters were cultured in the presence of WR99210 to induce structural stabilisation of the murine DHFR domain present in the reporter cargo [25]. Stabilisation of the mDHFR domain has been shown to improve cargo interaction with PTEX, enabling isolation of an otherwise transient interaction by co-immunoprecipitation (Fig 5A, panel 2)[38]. Parasites were then enriched by magnetic purification, lysed in 1% Triton X-100, and incubated with either anti-Nluc IgG, followed by antibody capture with protein A agarose, or anti-HA agarose beads (Fig 6A). Anti-Nluc immunoprecipitations of WT and P5 Lys Hyp1-Nluc-mDHFR-3xFLAG revealed that HSP101 interacted with WT Hyp1-Nluc-mDHFR-3xFLAG but not with the P5 Lys Hyp1-Nluc-mDHFR, likely due to stabilised interactions between WT Hyp1-Nluc-mDHFR-3xFLAG-3xFLAG and HSP101 in the PV where unfolding occurs (Fig 6B, lanes 13 and 16). To preserve weaker binding interactions, parasites were treated with 0.5 mM thiol-cleavable cross-linker dithiobis(succinimidyl propionate) (DSP) prior to lysis and immunoprecipitation. Under these conditions we found that HSP101 readily co-eluted with WT and P5 Lys Hyp1-Nluc-mDHFR-3xFLAG from parasites lysed in Triton X-100 (Fig 6B, lanes 14 and 17) and the more stringent RIPA buffer (Fig 6B, lanes 15 and 18). This data suggests that HSP101 can bind proteins destined for export in the ER prior to secretion into the PV. Given most of the immunoprecipitated P5 Lys Hyp1-Nluc-mDHFR-3xFLAG was not cleaved within the PEXEL motif (Fig 5D, S3 Fig), our data also suggests that correct PEXEL processing may not be necessary for HSP101 binding. To rigorously exclude the possibility that small amounts of PEXEL-cleaved P5 Lys Hyp1-Nluc-mDHFR-3xFLAG (Fig 5D and S3 Fig) was the species that co-immunoprecipitated HSP101, we also performed reciprocal anti-HA immunoprecipitation to capture the HA-tagged HSP101 (Fig 6C). Western blot analysis of the eluates indicated that HSP101-HA could co-immunoprecipitate both PEXEL-cleaved WT Hyp1-Nluc-mDHFR-3xFLAG (Fig 6C, lanes 11 and 13) and the non PEXEL-cleaved P5 Hyp1-Nluc-mDHFR-3xFLAG (Fig 6C, lanes 12 and 14). Furthermore, HSP101 did not co-immunoprecipitate SERA5, a protein secreted into the PV, validating the specificity of the interaction (Fig 6C).

**Fig 6.**
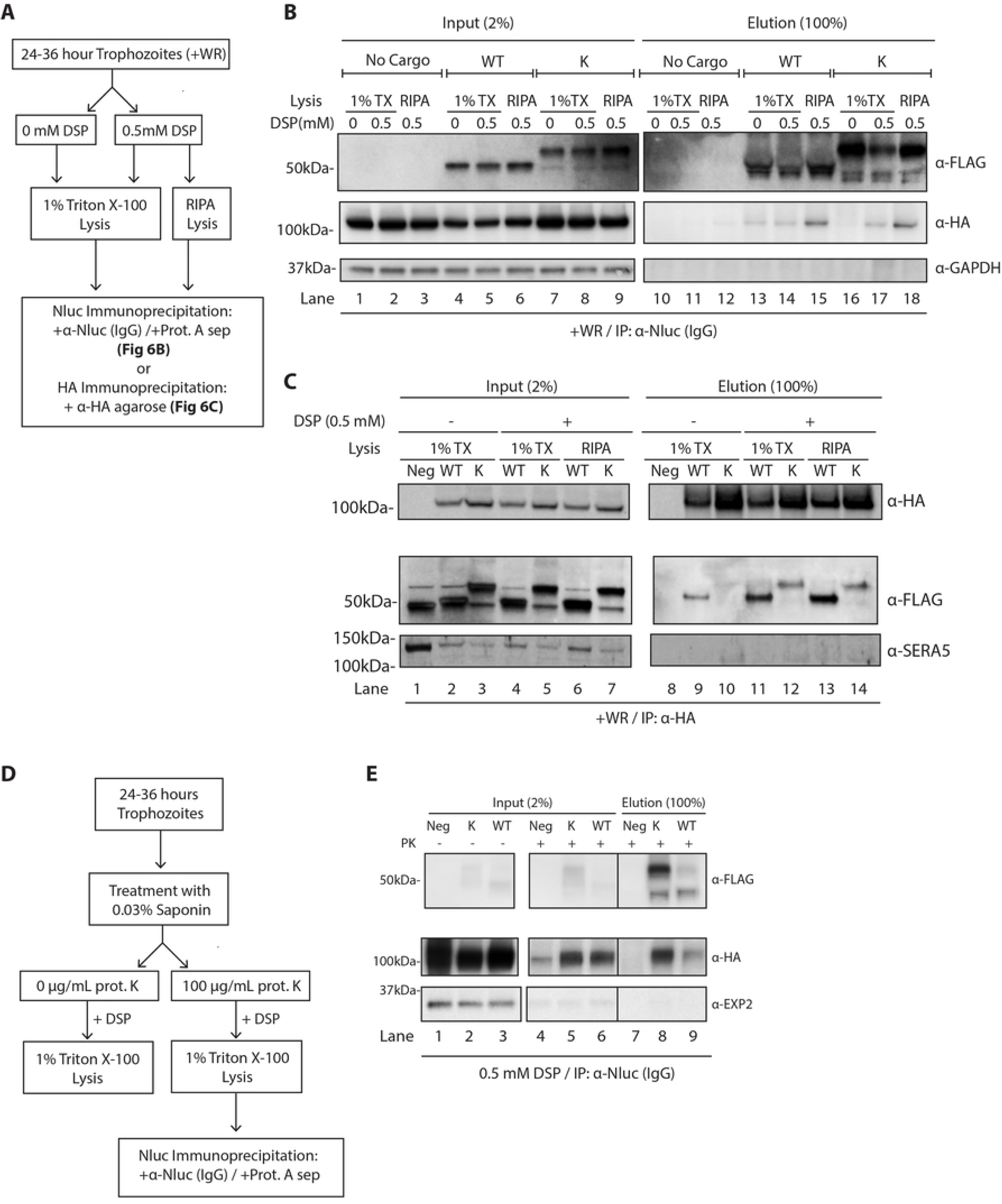
ER-located HSP101 interacts with Hyp1-Nluc-mDHFR-3xFLAG irrespective of Hyp1 PEXEL processing. (**A)** Schematic of parasite treatment and the subsequent co-immunoprecipitation to assess the interaction between Hyp1 reporters and HSP101. (B) Western blot of anti-Nluc immunoprecipitation performed with HSP101-HA*glms* parasites expressing WT or P5 Lys Hyp1-Nluc-mDHFR-3xFLAG. Where indicated, parasites samples were crosslinked with 0.5 mM DSP and lysed with either 1% TX-100 buffer or RIPA buffer. The data indicate that the mutant PEXEL reporter engages with HSP101 despite residing in the ER and not being cleaved by plasmepsin V. (C) Reciprocal anti-HA immunoprecipitation was performed using the same samples as (B) to confirm that HSP101 was interacting with both the PEXEL-cleaved (WT) and non PEXEL-cleaved (P5 Lys) Hyp1-mDHFR-3xFLAG reporters. Parasites that expressed Hyp1-mDHFR-3xFLAG without a HA-tagged version of HSP101 were used as a negative control (Neg). For both immunoprecipitations, inputs (2%) and eluates (100%) were fractionated by SDS-PAGE. (D) Schematic of parasite treatment with proteinase K and the subsequent co-immunoprecipitation experiment to examine if ER-resident HSP101 binds to Hyp1 reporter proteins. (E) Western blot analysis of co-immunoprecipitation of WT and P5 Lys Hyp1-mDHFR-3xFLAG and parental HSP101-HA*glmS* line (negative control, Neg) captured with anti-Nluc IgG. *In situ* crosslinking was performed using 0.5 mM DSP. Input (2%) and eluate (100%) were fractionated by SDS-PAGE. Immunoblots were stained with anti-FLAG antibody to visualise cargo, anti-HA to visualise HSP101 and anti-EXP2 to control for PVM disruption. The data indicate that both WT and P5 Lys Hyp1-mDHFR-3xFLAG bind to the proteolytically resistant ER-pool of HSP101.

To explore if other PTEX components could bind ER-trapped cargo, PTEX150 was immunoprecipitated and could only co-purify the WT Hyp1-Nluc-mDHFR-3xFLAG reporter and not the P5 Lys Hyp1-Nluc-mDHFR-3xFLAG, suggesting that only ER-resident HSP101 can form an interaction with the ER-trapped mutant cargo (S4 Fig).

To further demonstrate that HSP101-cargo interaction occurred in the parasite, HSP101-HA*glmS* parasites expressing WT and P5 Lys Hyp1-Nluc-mDHFR-3xFLAG reporters were first permeabilised by saponin and treated with 100 µg/mL proteinase K to degrade the contents of the host cell and the PV, including PTEX components and associated HSP101 (Fig 6D). As was observed previously, HSP101 was still present in all treated samples, while the PV-located EXP2 was almost fully degraded by proteinase K (Fig 6E, lanes 1-3 vs lanes 4-6). Following digestion, the resulting pellets were similarly treated with 0.5 mM DSP and solubilised with 1% Triton X-100 lysis buffer prior to anti-Nluc co-immunoprecipitation assay (Fig 6E). Analysis of the eluted fraction revealed that both WT and P5 Lys Hyp1-Nluc-mDHFR-3xFLAG co-immunoprecipitated HSP101 (Fig 6E, lanes 8 and 9). We observed less HSP101 co-eluted with WT Hyp1-Nluc-mDHFR-3xFLAG relative to the P5 Lys Hyp1-Nluc-mDHFR-3xFLAG probably because much of the WT Hyp1-Nluc-mDHFR-3xFLAG had already trafficked to the PV and/or into the host cell and would have been susceptible to proteinase K digestion after saponin treatment. Consequently, ER-localised P5 Lys Hyp1-Nluc-mDHFR-3xFLAG, which is protected from protease, co-purified greater amounts of HSP101.

Collectively, our immunoprecipitation data indicates that HSP101’s interactions with cargo happens intracellularly as well as in the PV.

### Internal HSP101 does not directly associate with plasmepsin V

Previous work found proteomic-based evidence for an interaction between HSP101 and plasmepsin V, the protease responsible for PEXEL cleavage [35], leading to a proposed model where plasmepsin V first cleaves the PEXEL motif of a protein destined for export in the ER and then passes the processed protein onto HSP101 to subsequently chaperone the protein to the PV for export [35, 36]. To determine if ER-resident HSP101 directly interacted with plasmepsin V to perform cargo handover, we first performed an anti-FLAG co-immunoprecipitation using HSP101-HA parasites expressing WT and P5 Lys Hyp1-Nluc-mDHFR-3xFLAG in the presence of 0.5 mM DSP crosslinker. We found that the P5 Lys Hyp1-Nluc-mDHFR-3xFLAG reporter co-precipitated both plasmepsin V and HSP101 whilst the exported WT Hyp1-Nluc-mDHFR-3xFLAG only co-precipitated HSP101 (Fig 7A). This suggests that the accumulation of P5 Lys Hyp1-Nluc-mDHFR-3xFLAG in the ER might stabilise HSP101-cargo-plasmepsin V interaction in the ER. Encouraged by this finding, we sought to perform the reciprocal co-immunoprecipitation on HSP101-HA parasites in the presence of P5 Lys Hyp1-Nluc-mDHFR-3xFLAG. The co-immunoprecipitation however, revealed that while HSP101-HA co-purified the P5 Lys Hyp1-Nluc-mDHFR-3xFLAG reporter as previously demonstrated (Fig 6C), plasmepsin V was conspicuously absent (Fig 7B). This data suggests that HSP101 and plasmepsin V may separately bind to proteins destined for export and therefore, HSP101 may not receive proteins directly from plasmepsin V after PEXEL cleavage.

**Fig 7.**
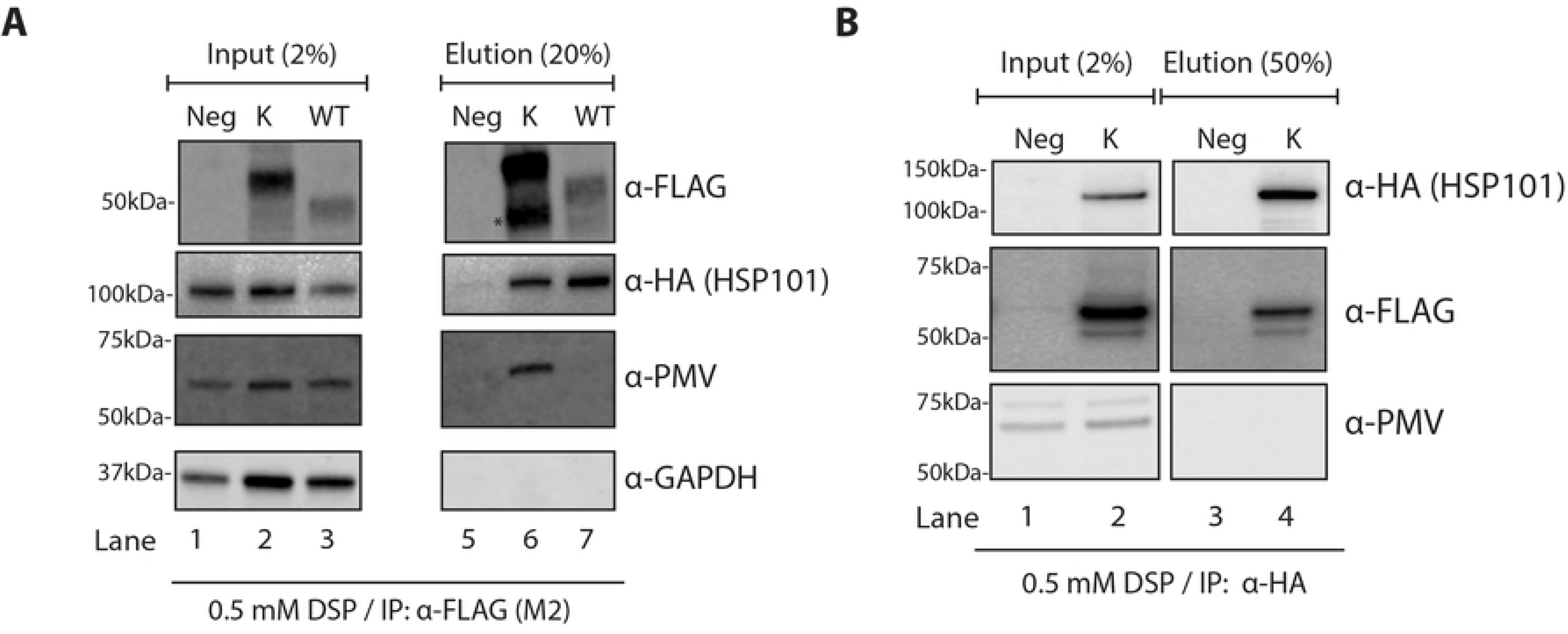
Plasmepsin V is not directly associated with HSP101 in the parasite ER. (A) Western blot analysis of WT and P5 Lys Hyp1-Nluc-mDHFR-3xFLAG (K) proteins captured with anti-FLAG M2 agarose beads indicate that the ER-trapped P5 Lys Hyp1-Nluc-mDHFR-3xFLAG associates with both HSP101 and plasmepsin V. Infected erythrocytes were magnetically purified then crosslinked with 0.5 mM DSP, lysed, and incubated with anti-FLAG M2 agarose beads. As a negative control, the HSP101-HA*glmS* parental parasites were treated similarly (Neg). Input (1.5%) and eluate (37.5%) were fractionated by SDS-PAGE. Immunoblot detection was performed using an anti-HA antibody to detect HSP101 as well as antibodies to plasmepsin V (PMV). Antibodies to GAPDH were used as a negative control. (B) Western blot of immunoprecipitated HSP101-HA from parasites expressing P5 Lys Hyp1-mDHFR-3xFLAG and probed with anti-plasmepsin V (PMV) indicate the protease is not directly associated with HSP101-HA. Input (2%) and eluate (50%) were fractionated by SDS-PAGE.

## Discussion

The trafficking route that targets PEXEL proteins post plasmepsin V cleavage from the ER to the PV has been enigmatic. Following arrival at the PV, it has not been clear how PEXEL proteins are selected for export by the PTEX from other PV-resident proteins [14, 50]. We have now presented three key findings to suggest that HSP101 may be the key component that mediates these two processes.

Firstly, our BN-PAGE and co-immunoprecipitation data suggest that EXP2 and PTEX150 form a stable subcomplex in the absence of HSP101. EXP2 or PTEX150 alone cannot interact with HSP101, suggesting that EXP2 has to associate with PTEX150, presumably to drive molecular changes that functionalise the subcomplex into a form permissive for HSP101 binding. The other implication of this finding is that HSP101 may therefore be able to dock on and off the EXP2/PTEX150 subcomplex without disrupting it. The ability to assemble and disassemble from the membrane-associated EXP2/PTEX150 subcomplex may be integral for HSP101 function, especially given that HSP101 was shown to retain its ability to associate with exported proteins when dissociated from EXP2 and PTEX150 [32]. For example, free HSP101 would be able to associate with other PV-resident proteins to retrieve cargo proteins [44], extract transmembrane proteins from the plasma membrane [31], or in the current context, enable cargo-bound HSP101 from the ER to reconstitute the full translocon.

This leads to our second key finding. We were able to show that *P. falciparum* HSP101 resides in the parasite’s ER in addition to its vacuolar localisation. Our data indicate that the parasite’s total pool of HSP101 appears to be split between the PV and ER and that the ER-located HSP101 is continuously secreted into the PV even during the timeframe of low transcription and protein synthesis of HSP101 [42]. Combined with the observation that five hours of BFA treatment at mid trophozoite stage depleted PV HSP101, our data suggest that much of ER-resident HSP101 may of PV origin rather than newly synthesised HSP101 and presumably the chaperone cycles between these two compartments. As a counterpoint, we cannot completely discount the possibility that the steady state levels of HSP101 are maintained through continuous secretion of newly made HSP101 and the degradation of PV HSP101. Given that PTEX components have been shown to undergo relatively little protein turnover throughout the entire 48-hour asexual cycle [27], we favour the former scenario.

Thirdly, we have provided evidence that the ER HSP101 is likely to bind specifically to PEXEL proteins *en route* to the PV. This is based on our finding that the ER HSP101 engages with our Hyp1 reporter protein in the ER and not secreted PV-resident protein SERA5. HSP101-cargo interaction in the ER occurs relatively transiently and does not seem to be stabilised by the unfolding of cargo. The mechanistic reason for this remains to be investigated, however, ClpB-like chaperones are known to transiently engage cargo when bound to ATP before committing to high affinity unfolding that is driven by the hydrolysis of bound ATP molecules [51, 52]. Therefore, it is possible that the ER environment only allows the formation of an initial weak interaction of cargo with HSP101, perhaps to prevent premature unfolding. Whilst we anticipated that the formation of the cleaved and acetylated PEXEL N-terminus might be important for HSP101 binding in the ER, we were surprised to find that HSP101 interacts with the P5 Lys Hyp1-Nluc-mDHFR-3xFLAG reporter which was not efficiently cleaved within the PEXEL motif by plasmepsin V. Although this might not be biologically relevant, it is tempting to speculate that perhaps the Ac-xE/Q/D motif of the mature PEXEL motif is not the binding site for HSP101.

Based on our findings, we propose that given HSP101’s affinity for cargo proteins in the ER and for the PTEX150-EXP2 subcomplex in the PV, this provide the means by which exported proteins are selectively translocated upon arrival at the PV. In other words, only proteins that can bind to HSP101 in the ER, can get access to the PTEX150-EXP2 subcomplex upon arrival at the PV, with binding of HSP101 leading to assembly of the full PTEX complex and the stimulation of protein translocation (Fig 8). PV-resident proteins do not interact with HSP101 in the ER and thus do not have access to the translocon once they arrive at the PV (Fig 8).

**Fig 8.**
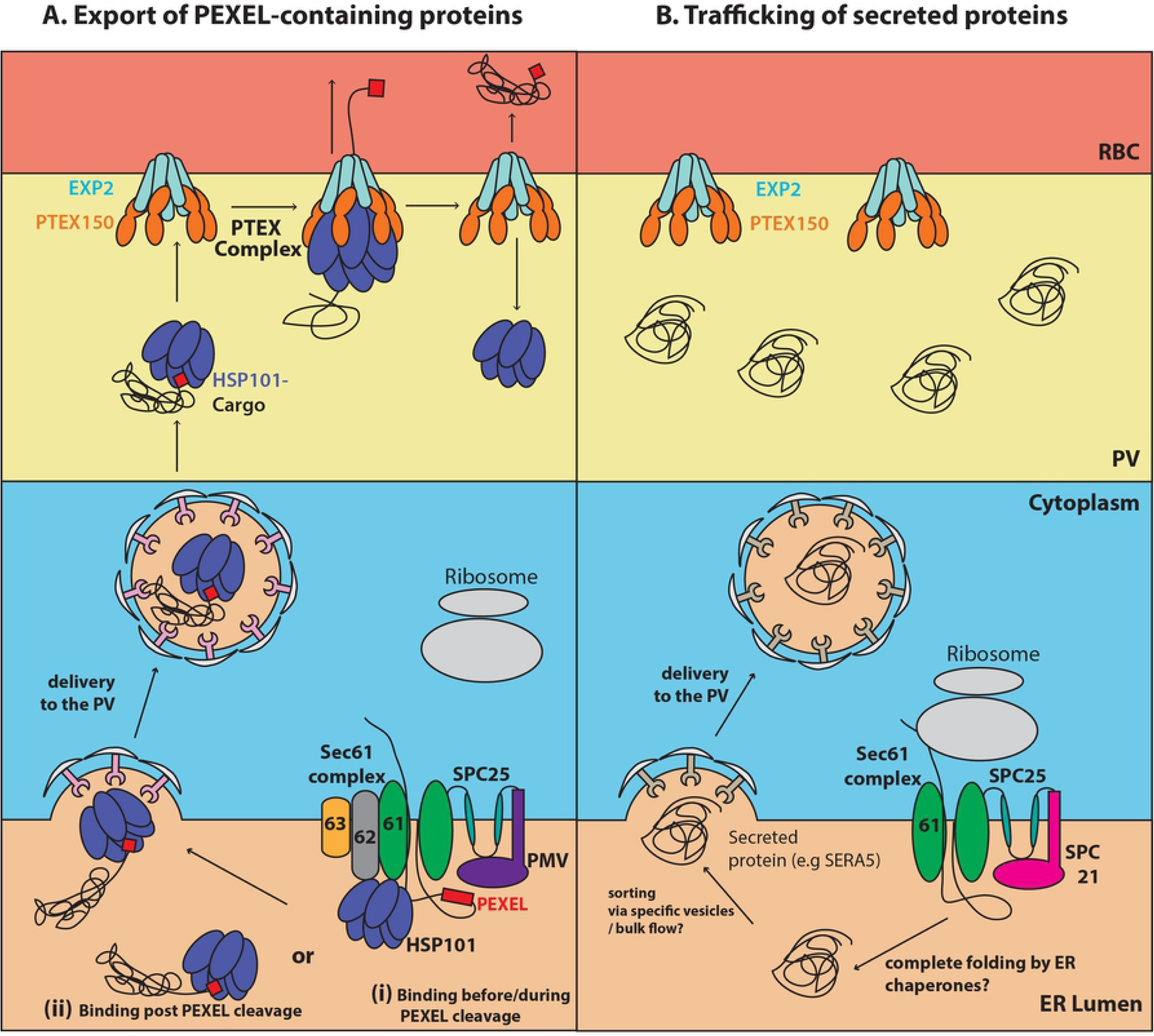
Model of PEXEL protein targeting to the vacuolar translocon by HSP101. (A) PEXEL-containing proteins are imported after translation into the ER via a Sec61/62/63-SPC25-PMV translocon [23] and are recognised by HSP101, either during import (i) or after PEXEL processing (ii), prior to their release into the ER lumen. HSP101-bound cargo is then trafficked to the parasite membrane via the vesicular trafficking system and released into the PV. HSP101 affinity for the PTEX150-EXP2 subcomplex at the vacuolar membrane drives reconstitution of the full PTEX complex capable of translocating the cargo into the host cell. The model depicted here is described for soluble PEXEL proteins, but it is possible that the same mechanism governs the trafficking of transmembrane proteins. (B) Proteins with a classical N-terminal signal sequence destined for the PV such as SERA5, are imported into the ER by the Sec61-SPC25-SPC21 complex [23]. After signal sequence cleavage, the released proteins are folded and secreted into the PV. Since these proteins are not bound to HSP101, they cannot translocate via PTEX.

Several questions remain to be addressed. For instance, the exact sequence requirement for cargo recognition by HSP101 is currently unknown but residues immediately downstream of the PEXEL motif have been shown to be important for export and are functionally equivalent to the N-termini of PNEPs [53, 54]. If HSP101 specificity is conferred by this downstream region, rather than by the presence of cleaved mature PEXEL motif, it would explain the ability of HSP101 to recognise both PEXEL and PNEPs [55]. Further analysis of what information is contained within the region downstream of the PEXEL motif and what features might unify it with regions of exported PNEPs should therefore be a priority, and in particular to search for features beyond simple primary sequence similarities [50].

The precise timing of cargo recruitment by HSP101 and how it is linked to plasmepsin V cleavage that licenses the proteins for export is also not clear at this point. It has been hypothesised that plasmepsin V could hand over PEXEL proteins post cleavage to HSP101, that in turn chaperones the cargoes to the PV via the classical secretory pathway [35]. Our data showed that there is no direct association between HSP101 and plasmepsin V although both were co-purified with our ER-trapped P5 Lys Hyp1 reporter. This could mean that the cleaved cargo is passed to one or more intermediate factors post plasmepsin V cleavage before being taken by HSP101, or alternatively, HSP101 competes with plasmepsin V for cargo binding. A alternative explanation could also be proposed, where HSP101 binds to cargo prior to plasmepsin V cleavage, perhaps through interaction with the ER import translocon [23, 56], and releases it momentarily for plasmepsin V to cleave the PEXEL motif before immediately resuming its interaction with the mature cargo.

Another important question is what prevents the cargo from being translocated through the central cavity of the HSP101 hexamer once it engages with cargo in the ER, prior to binding the rest of PTEX? It has been proposed that at the PVM, PTEX150 could promoting HSP101 unfolding activity through stimulate HSP101’s unfoldase activity after engagement with the PTEX150-EXP2 subcomplex [50]. The absence of PTEX150 from the ER would prevent HSP101 from unfolding the cargo prematurely until the HSP101/cargo complex reached the rest of PTEX in the PV. Although our model (Fig 8) still requires further validation, our findings have revised our current understanding of protein export and put forward a new and testable model for future research.

## Materials and Methods

### Molecular cloning

The expression of Hyp1-Nluc-mDHFR-3XFLAG reporters was driven by a bidirectional *Plasmodium berghei* EF1a promoter that also controlled expression of the blasticidin deaminase drug resistance cassette. The plasmid pEF-Hyp1-Nluc-mDHFR-3xFLAG was derived from plasmid pEF-Hyp1-Nluc-DH-APEX [38]. The Hyp1 component of this plasmid contained the first 113 aa of Hyp1 (PF3D7_0113300), including the RLLTE PEXEL motif [46] which when cleaved leaves 52 aa of Hyp1 remaining. A synthetic murine dihydrofolate reductase (mDHFR) gene fragment with C-terminal 3x FLAG epitopes (Bioneer Pacific) was ligated into the Nluc-DH-APEX plasmid using *SpeI* and *MluI* enzymes to remove the previous mDHFR-APEX gene cassette. This generated the final pEF-Hyp1-Nluc-mDHFR-3xFLAG plasmid.

Generation of the P5 lysine (lys/K) mutation of the Hyp1-Nluc-mDHFR-3xFLAG was performed as follows: Hyp1 region was first amplified as two overlapping PCR fragments. First fragment amplified with Hyp1K_1 (5’-TGCTTATAAATAAATAAAAATTTTATAAAA<u>CTCGAG</u>CAAAATGA-3’) and Hyp1K_2 (5’-TGCTAGCAGGTTATTAACAAAATATAAAGACACATTACA-3’) and the second fragment amplified with Hyp1K_3 (5’-TGTAATGTGTCTTTATATTTTGTTAATAACCTGCTAGCA-3’) and Hyp1K_4 (5’-TCGAGTGTGAAGA<u>CCATGG</u>TATCAACAACA-3’).

The overlap region between these two PCR products contained the lysine mutation indicated above. PCR fragments were then sewn together with primer pair Hyp1K_1 & Hyp1K_4 via overlapping PCR, ligated into the pJET1.2/blunt plasmid (ThermoFisher Scientific), and electroporated into *E. coli* XL-10 gold cells (Stratagene). After sequencing, the mutant P5 Lys Hyp1 fragments were released from pJET1.2/blunt using *XhoI* and *NcoI* and ligated into pEF-Hyp1-Nluc-mDHFR-3xFLAG to replace the wildtype Hyp1 fragment. This resulted in the final pEF-P5Lys Hyp1-Nluc-mDHFR-3xFLAG plasmid.

To generate plasmid pHSP101-HA*glmS* for creating a *P. falciparum* HSP101 conditional knockdown parasite line, the parent construct pPTEX150-HA-glmS [57] was digested with *Bg*lII and *Pst*I to remove the *Pfptex150* targeting sequence and in its place, 920 bp of the C-terminus of *Pfhsp101* sequence minus the stop codon was cloned. This *Pfhsp101* targeting sequence was amplified from *P. falciparum* 3D7 gDNA using oligonucleotides DO610 (5’-GCG<u>AGATCT</u>CTAAATCCATTATTGGAAATGAAGAT-3’) and DO611 (5’-GCG<u>CTGCAG</u>CGGTCTTAGATAAGTTTATAACCAAG-3’). PCR genotyping was performed by amplifying parasite genomic region with a primer set DO561 (5’-TAAGTACATTAGAAAAGGATGTAGACA-3’) / DO562 (5’-GGTCTTAGATAAGTTT-ATAACCAAGTT-3’), DO561 / DO233 (5’-tccgcatgcGGTCTTAGATAAGT-TTATAACCAAGTT-3’), and DO561 / DO276 (5’-GTGATTTCTCTTTGTTCAAGGA-3’) as indicated in Fig 1A,B.

### Parasite maintenance and generation of transgenic parasites

Asexual blood stage *Plasmodium falciparum* was cultured as per [58]. Cultures were maintained routinely in complete media comprising of RPMI-1640 base media supplemented with 2.5mM HEPES, 367µM Hypoxanthine, 31.25µg/mL Gentamicin, 25mM NaHCO_3_, and 0.5% Albumax II (Invitrogen). To generate HSP101-HA*glmS* parasites, 100 µg of pHSP101-HA*glmS* plasmid was electroporated into erythrocytes which were then mixed with 3D7 trophozoites and selected with 2.5 nM WR99210 [59, 60] Parasites were cycled on and off WR99210 until a 100 kDa HA positive band appeared by western blot using anti-HA antibodies, indicative of integration into the *hsp101* locus. Clonal HSP101-HA*glmS* positive parasites were selected by limiting dilution and correct integration verified by PCR.

To induce episomal expression of Hyp1-Nluc-mDHFR-3xFLAG and P5 Lys Hyp1-Nluc-mDHFR-3xFLAG in HSP101-HA*glmS* parasite line, 100 µg of pEF-Hyp1-Nluc-mDHFR-3xFLAG and pEF-P5 Lys Hyp1-Nluc-mDHFR-3xFLAG was used to transfect the HSP101-HA*glmS* parasite line. Transfections were performed as above except that 2.5 µg/mL blasticidin S was used for selection. Cargo trapping was performed essentially as described before [38] with young ring stage parasites treated with 10 nM WR99120 for 16-20 hours before being harvested for immunoprecipitation or IFA (described below).

### PTEX knockdown and Blue Native PAGE

Saponin lysed parasite pellets were solubilized in 1% digitonin (ThermoFisher) for 30 min at 4°C. Following incubation, the soluble component was separated from the insoluble pellet by centrifugation at 14,000 x g for 30 min. 5% Coomassie G-250 and 100% glycerol was spiked into the supernatant to reach final concentrations of 0.25% and 10% respectively. Samples were separated on a 3-12% NativePAGE Bis-Tris gel (ThermoFisher). Gels were run as per manufacturer prior to blotting onto a 0.2 µm polyvinylidene fluoride (PVDF) membrane using iBlot® system (Invitrogen). To enable the detection of HSP101, the gel was incubated for 15 minutes at room temperature in 50 mM DTT and 0.1 % SDS in the tris-glycine buffer prior to transfer. The membranes were subsequently fixed with 8% (v/v) acetic acid in water, air-dried, and incubated with 100% methanol to remove the excess Coomassie G-250 dye. The membrane was blocked in 1% casein in 1x PBS for 1 h at room temperature prior to probing with primary antibody solutions (diluted in 1% casein in PBS) as following: rabbit anti-HSP101 (IgG purified; 30 µg/mL), rabbit anti-PTEX150 (r741, 1:500), mouse anti-EXP2 (10 µg/mL).

### Indirect immunofluorescence analysis

IFA was performed essentially according to [61]. Where iRBCs were settled onto a poly-L-lysine (Sigma, P8920) coated coverslip and fixed with 4% paraformaldehyde/0.0025% glutaraldehyde. Following fixation, the cells were permeabilised with 0.1 M glycine/0.1 % Triton X-100 for 12 minutes at RT. Alternatively, thin blood smears of parasites were fixed in 90% acetone/10% methanol for 5 mins where specified. Coverslips or slides were blocked with 3 % BSA/0.02 % Triton X-100/1x PBS and probed overnight at 4°C, with rabbit anti-Nluc (12.5 µg/mL), mouse anti-EXP2 (10 µg/mL), rabbit anti-ERC (1:1000), mouse anti-FLAG M2 (Sigma, 10 µg/mL), mouse anti-HA (Sigma clone HA-7; 1:500), rabbit anti-SBP1 (1:500) [33] and rabbit anti-STEVOR (PF3D7_1254100, 1:500) [62]. After washing goat anti-rabbit Alexa Fluor 594 and Goat anti-mouse Alexa Fluor 488 (1:2000) secondary antibodies were applied for one hour at RT. Fixed material was mounted in VECTASHIELD with DAPI and imaged on Zeiss Cell Axio Observer (Carl Zeiss). Image acquisition was performed with Zen Blue imaging software.

### Quantification of internal HSP101 staining

Analysis was performed essentially according to the “*in vitro* cyst DBA fluorescence quantification” protocol [63]. Images were first imported to the open java source ImageJ/FIJI for analysis and “set measurement” function was set to record area, integrated intensity, and mean grey values. Firstly, a region of interest (ROI) was first drawn around the parasite cell (including the PV) using the freehand selection tool. DIC channel and the anti-HA/anti-EXP2 staining was used concomitantly as a guide to define the outermost part of the PV. Measurement data for total parasite fluorescence intensity (TPFI) were then recorded. An identical procedure was carried out to measure a ROI inside the parasite cell excluding the PV (Internal fluorescence, IF). The whole procedure was repeated 3 times for each cell to minimize errors. The background intensity was measured inside 3 uninfected erythrocytes within each captured field and was averaged to obtain mean background value (MBV). Total parasite and internal fluorescence intensity was corrected by subtracting the MBV from the averaged integrated density of TPFI or IF with the value of MBV multiplied by the mean area of the measured cell. The estimated PV fluorescence intensity for each cell was calculated by subtracting the cell’s corrected TPFI and IF integrated density values. Lastly, the calculated intensity for PV was divided by the IF value to calculate the amount of internal fluorescence intensity relative to the PV. The internal/PV ratios of >10 cells were measured for each category and plotted on GraphPad Prism 8.2 software to calculate the statistical significance by ordinary one-way ANOVA test.

### 3D-Structured Illumination Microscopy (3D-SIM)

3D-structured illumination microscopy (3D-SIM) was performed using a Nikon Ti-E inverted microscope (Nikon) equipped with a motorised piezo stage, Nikon Intensilight E, SIM illuminator, SIM microscope enclosure, and Perfect Focus System. Cells were prepared as per IFA protocol but then imaged using a Plan Apo VC 100x 1.4NA oil objective and using 2D/3D-SIM imaging modes with diffraction grating 3D 1 Layer for 2D/3D SIM. A Z-stack for each cell was taken for co-localisation analysis using the open source java ImageJ’s JaCoP plugin.

### Biochemical PEXEL cleavage assays

The cleavage assay was performed essentially as described in Hodder *et al* [64]. 2 nM of *P. vivax* plasmepsin V in buffer (25 mM Tris-HCl and 25 mM MES, pH 6.4) was incubated with 5 μM FRET peptide substrates representing WT and mutant KAHRP (DABCYL-RNK**RTLAQ**KQ-E-EDANS and DABCYL-RNK**ATAAQ**KQ-E-EDANS) and Hyp1 (DABCYL-G-KI**RLLTE**YKD-E-EDANS and DABCYL-G-KI**RLLTK**YKD-E-EDANS) sequences in a total volume of 20 μL. Samples were incubated at 20°C for 20 h and were measured with an Envision plate (PerkinElmer) reader (ex. 340 nm; em. 490 nm). Raw fluorescence values were subtracted from fluorescence background values and then data for each substrate was normalized to 100% activity. Biochemical *P. vivax* plasmepsin V inhibitory assays (20 μL total volume) consisted of 2 nM of *P. vivax* plasmepsin V in buffer (25 mM Tris-HCl and 25 mM MES, pH 6.4) with 5 μM FRET fluorogenic peptides. Inhibitory values were determined with a nonlinear regression four-parameter fit analysis in which the parameters were not constrained using Domatics software (version 5.3.1612.8630).

### Immunoblotting analysis

Protein samples were electrophoresed on a NuPAGE 4-12% Bis-Tris gel (Invitrogen) and transferred onto a nitrocellulose membrane using iBlot® system (Invitrogen). The membrane was subsequently blocked using 1% casein in 1x PBS and incubated with primary antibody solutions with following dilutions: mouse anti-HA (Sigma, 1:500), rabbit anti-HSP101 (r950, 1:500), rabbit anti-PTEX150 (r741, 1:500), mouse anti-EXP2 (5-10 µg/mL), rabbit anti-EXP2 (r1167, 1:1000), chicken anti-FLAG (Abcam,1:2000), mouse anti-FLAG (10 µg/mL), rabbit anti-Nluc (12.5 µg/mL), rabbit anti-plasmepsin V (1:2000), rabbit anti-GAPDH (1:2000), rabbit anti-SERA5 (1:1000) for 1 hour at RT or overnight at 4°C. Secondary antibodies were incubated for 1-2 hours in following dilutions: Goat anti-rabbit Alexa Fluor Plus 700 and 800, Goat anti-mouse Alexa Fluor Plus 700 and 800 (Invitrogen 1: 10,000), and horseradish peroxidase-conjugated Goat anti-chicken IgY (Abcam, 1: 10,000). For enhanced chemiluminescence detection, membranes were probed with SuperSignal^TM^ West Pico PLUS substrate (Pierce). All imaging and densitometry analysis were performed using Odyssey Fc system (LI-COR).

### Protease protection assays

Protease protection assays were completed as outlined in [30]. Briefly, magnet purified trophozoites (24-30 hpi) were pelleted (500x*g*/ 5 min) and washed twice with digestion buffer 50 mM Tris, 150 mM NaCl, 1 mM CaCl_2_, pH 7.4. The resultant pellet was split into 12 equal fractions. Four tubes were subsequently treated with recombinant equinatoxin made in house and empirically determined to lyse ~100% of erythrocytes. Another four tubes were treated with equinatoxin and 0.03% saponin; and the remaining four tubes were treated with equinatoxin and 0.25% Triton X-100. Parasites were incubated at room temperature for 10 min/ shaking at 1000 rpm. Proteinase K was then added to selected tubes to final concentrations of 0, 1, 5 and 20 μg/mL and digested for 15 min at 37°C. The reaction was stopped by addition of 200 μL of cOmplete^TM^ Protease Inhibitor Cocktail (Roche) made up at a high concentration of 2 tablets per 3 mL digestion buffer with 1 mM PMSF. Parasite material was centrifuged (1000 x *g*, 5 min) and pellet material was separated from supernatant. Pellet and supernatant material was resuspended in sample buffer and electrophoresed via NuPAGE Bis Tris SDS-PAGE 4-12% prior to analysis by western blotting.

### Co-immunoprecipitation assays

For HA co-immunoprecipitations, trophozoite iRBCs were treated with 0.09% saponin to remove haemoglobin and solubilised in 20x cell pellet volume of lysis buffer (1% Triton X-100, 0.1% SDS, 150 mM NaCl, 10 mM Tris-HCl pH 7.4) supplemented with cOmplete^TM^ Protease Inhibitor Cocktail (Roche) and subjected to 2 freeze and thaw cycles. The lysate was clarified by centrifugation (16,000 x *g*, 10 minutes, 4°C), and total protein concentration was measured by Bradford assay. Lysates were adjusted to 1-2 mg total protein in 1 mL and incubated overnight at 4°C with 25 µL of packed Monoclonal Anti-HA Agarose (Sigma-Aldrich). Following incubation, agarose beads were washed 5x with 1 mL the lysis buffer and eluted with 50 µL 2x protein sample buffer (100 mM Tris-HCl pH 6.8, 4 mM EDTA, 4% SDS, 0.01% bromophenol blue, 20% (v/v) glycerol).

For Hyp1-Nluc-mDHFR-3xFLAG co-immunoprecipitations, trophozoite iRBCs were magnetically purified and solubilised as above. Lysates were adjusted to 0.5-1 mg total protein in 1mL and incubated overnight at 4°C with 10 µg IgG-purified anti-Nluc antibody. Following incubation, 25 µL packed Recombinant Protein A-Sepharose 4B (Invitrogen) was used to capture the immune complexes for 1 hour at RT. Beads were subsequently washed 5x with 1 mL lysis buffer and eluted with 50 µL protein sample buffer.

FLAG immunoprecipitations, iRBCs were magnetically captured and were solubilised in 20x pellet volume of 0.5x RIPA buffer (0.5% Triton X-100, 0.5% sodium deoxycholate, 0.05% SDS, 25 mM Tris pH 7.4, 150 mM NaCl). The rest of the immunoprecipitations was performed as above except that anti-FLAG M2 agarose beads (Sigma-Aldrich) were used to capture the reporter protein and all washes were performed in RIPA buffer.

For co-immumoprecipitations with PTEX antibodies, saponin lysed late ring-stage parasites that had been cross-linked with 1 mM DSP were solubilised with 0.5% Triton X-100 in 1x PBS supplemented with cOmplete^TM^ protease inhibitor cocktail (Roche) at 4°C. Clarified lysates were incubated overnight at 4°C with 10 µg rabbit anti-HSP101 IgG or 20 µL of rabbit polyclonal anti-PTEX150 serum [26]. Following overnight incubation, 50 µL packed protein A-Sepharose 4B (Invitrogen) was added to the immune complexes and samples were incubated for an additional 1 hour at RT. Beads were subsequently washed 5x with 1 mL IP 0.5% Triton X-100 in 1x PBS and eluted with 60 µL of protein sample buffer.

### Mass spectrometry

Coomassie stained protein bands were excised and destained with 50% acetonitrile (ACN) in 100 mM ammonium bicarbonate (ABC) pH 8.5. The proteins were reduced and alkylated after treatment with 10 mM DTT (Astral Scientific) and 20 mM chloroacetamide (Sigma). The gel was dehydrated using 100% ACN and rehydrated with digestion solution, containing 100ng/μL trypsin (Promega) or 100ng/μL GluC (Promega) in 100 mM ABC. After overnight digestion at 37°C, the tryptic peptides were extracted from the gel and subject LC–MS/MS analysis.

Enzyme digests were analysed by LC-MS/MS using the QExactivePlus mass spectrometer (Thermo Scientific, Bremen, Germany) coupled online with an Ultimate 3000 RSLCnano system (Thermo Scientific, Bremen, Germany). Samples were concentrated on an Acclaim PepMap 100 (100 μm × 2 cm, nanoViper, C18, 5 μm, 100 Å; Thermo Scientific) trap column and separated on an Acclaim PepMap RSLC (75 μm × 50 cm, nanoViper, C18, 2 μm, 100 Å; Thermo Scientific) analytical column by increasing concentrations of 80% acetonitrile/0.1% formic acid at a flow of 250 nL/min for 90 min.

The mass spectrometer was operated in the data-dependent acquisition mode to automatically switch between full scan MS and MS/MS acquisition. Each survey full scan (m/z 375–1575) was acquired in the Orbitrap with 60,000 resolution (at m/z 200) after accumulation of ions to a 3 × 106 target value with maximum injection time of 54 ms. Dynamic exclusion was set to 15 s to minimize repeated selection of precursor ions for fragmentation. The 12 most intense multiply charged ions (z ≥ 2) were sequentially isolated and fragmented in the collision cell by higher-energy collisional dissociation (HCD) with a fixed injection time of 54 ms, 30,000 resolution and automatic gain control (AGC) target of 2 × 105. The raw files were analysed Proteome Discoverer v2.5 (Thermo Scientific) and searched against a custom database of the recombinant sequence appended to the *Plasmodium falciparum* UniProtKB using the Byonic v3.0.0 (ProteinMetrics) search engine to obtain sequence information. Only proteins falling within a predefined false discovery rate (FDR) of 1% based on a decoy database were considered further.

## Acknowledgements

The authors would like to thank Brian Cooke for anti-SBP1 antibody, Michael Duffy and Michaela Petter for anti-STEVOR antibody, Leanne Tilley and Matthew Dixon for the anti-ERC and anti-GAPDH antibodies, Alan Cowman and Danuskha Marapana for the anti-plasmepsin V antibody and recombinant *P. vivax* plasmepsin V and WEHI monoclonal antibody facility for generating anti-FLAG and anti-EXP2 IgGs used in this study. We thank Nghi Nguyen, Anna Ngo, Chad Johnson, Paul Sanders, and Benjamin Dickerman for their technical assistance and the Australian Red Cross Blood Bank for providing human blood and Jacobus Pharmaceutical for WR99210.

## Supporting Information

**S1 Fig. Knockdown of HSP101 does not disrupt the interaction of EXP2 and PTEX150 subcomplex.**

(A) BN-PAGE analysis of EXP2, PTEX150, and HSP101-HA*glmS* parasite lines following glucosamine treatment. Saponin-lysed pellets were lysed with 1% digitonin, separated on 4-12% NativePAGE gel, and analysed via western blotting using monoclonal anti-HA antibody. The 1236 kDa PTEX bands and the major oligomeric species of EXP2, PTEX150, and HSP101 seen with PTEX-specific antibodies were also recognised by the anti-HA antibody (red text; see Fig 2A, 2B, and 2C). Additional HA-specific bands were also observed in the case of EXP2 and PTEX150-HA*glmS* (black text). In the case of HSP101-HA*glmS*, only the 1236 kDa and a faint >720 kDa band were observed (black text), indicating that the 1048 kDa and the 200 kDa bands found with anti-HSP101 antibody (Fig 2A) were likely non-specific. (B) Representative western blots of PTEX co-immunoprecipitated eluates to establish which PTEX components could still form subcomplexes when another component was knocked down (n=3 independent biological replicates). Parasites expressing HA-*glmS* tagged PTEX core components were treated +/− 2.5 mM glucosamine (GlcN) and immunoprecipitation (IP) was performed using either anti-PTEX150 antibodies (for HSP101 and EXP2 knockdown) or anti-HSP101 antibodies (for PTEX150 knockdown). (C) Quantification of PTEX band intensity represented in (A) where the amounts of PTEX proteins were normalised to the immunoprecipitated protein. Knockdown (Kd) of HSP101 did not appear to block the interaction between PTEX150 and EXP2. Knockdown of EXP2, however, disrupts PTEX150’s interaction with HSP101. The knockdown of PTEX150 was only 20% and thus HSP101’s interaction with EXP2 was therefore not greatly reduced. Plotted data represents the mean ±SD (n=3 independent replicates). Statistical significances were measured using an unpaired t-test with Welch’s correction. p-values are indicated on the graph.

**S2 Fig. P5 Lys Hyp1-Nluc-mDHFR-3xFLAG possesses a poorly cleavable PEXEL motif.**

5 µM fluorogenic peptides were incubated with 2 nM recombinant *P. vivax* PMV and assayed at 20°C. Fluorescence data was normalised to the WT substrates (n=3, error bars = SD) and indicated that the cleavage of P5 Lys Hyp1 peptide (RLLTK) was inhibited to the same level as the double P1 and P3 KAHRP mutant peptide (RTLAQ to ATALQ). Statistical significance was determined using ordinary one-way ANOVA. ****, p-value<0.0001, ***, p-value<0.001.

**S3 Fig. Proteomic analysis of immunoprecipitated WT and P5 Lys Hyp1-Nluc-mDHFR-3xFLAG reporter proteins indicated the mutant protein was not cleaved within the PEXEL motif.**

(A) Proteins immunoprecipitated using anti-FLAG IgG beads from parasites expressing WT or P5 Lys Hyp1-Nluc-mDHFR-3xFL reporter proteins were fractionated by SDS-PAGE. Protein bands were visualised with Coomassie stain and protein bands corresponding to the molecular weight of the PEXEL-cleaved WT Hyp1 (band 2) and the miscleaved P5 Lys (band 3) were excised (red boxes). The matching region of the gel for WT (band 1) and P5 Lys (band 4) were also excised and subjected to the same analysis. The protein bands were digested with trypsin or GluC and subjected to mass spectrometry to identify peptide fragments. (B) The amino acid sequence of WT Hyp1 region of Hyp1-Nluc-mDHFR-3xFL protein showing PEXEL motif (underlined, red), peptide cleavage site (arrow), transmembrane domain (blue). Below this is a diagram of the full-length reporter protein with peptide coverage of protein bands 1 and 2 (B1 and B2) indicated in green. (C) Peptide coverage of P5 Lys Hyp1-Nluc-mDHFR-3xFL reporter protein bands 3 and 4 (B3 and B4) as described for (B). Peptides identified between the transmembrane domain and PEXEL motif indicate the mutant protein is processed upstream of the PEXEL motif.

**S4 Fig. PTEX150 interacts with the exported WT Hyp1 reporter but not with the ER-trapped P5 Lys Hyp1 reporter.**

PTEX150 was immunoprecipitated from HSP101-HA*glms* parasites expressing WT or P5 Lys Hyp1-Nluc-mDHFR-3xFLAG reporters using polyclonal anti-PTEX150 (r942; against the C-terminal region of PTEX150). The parasites were either lysed with 1% TX-100 buffer or RIPA buffer or were crosslinked with 0.5 mM DSP and input (2%) and eluates (100%) were fractionated by SDS-PAGE. Western blots indicate that the WT Hyp1-Nluc-mDHFR-3xFLAG reporter interacts with PTEX150 as part of the PTEX complex with EXP2 and HSP101 but not with the ER-trapped P5 Lys Hyp1-Nluc-mDHFR-3xFLAG reporter.

